# Deficiency in endocannabinoid synthase *DAGLB* contributes to Parkinson’s disease and dopaminergic neuron dysfunction

**DOI:** 10.1101/2021.12.09.471983

**Authors:** Zhenhua Liu, Nannan Yang, Jie Dong, Wotu Tian, Lisa Chang, Jinghong Ma, Jifeng Guo, Jieqiong Tan, Ao Dong, Kaikai He, Jingheng Zhou, Resat Cinar, Junbing Wu, Armando Salinas, Lixin Sun, Justin Kung, Chengsong Xie, Braden Oldham, Mantosh Kumar, Sarah Hawes, Jinhui Ding, Lupeng Wang, Tao Wang, Piu Chan, Zhuohua Zhang, Weidong Le, Shengdi Chen, David M. Lovinger, Guohong Cui, Yulong Li, Huaibin Cai, Beisha Tang

**Affiliations:** Transgenic Section, Laboratory of Neurogenetics, National Institute on Aging, National Institutes of Health, Bethesda, MD 20892, U.S.A.; Department of Neurology, Xiangya Hospital, Central South University, Changsha, Hunan 410008, China; Clinical Research Center on Neurological Diseases, the First Affiliated Hospital, Dalian Medical University, Dalian, Liaoning, 116011, China; Department of Neurology, Ruijin Hospital Affiliated to Shanghai Jiao Tong University School of Medicine, Shanghai 20025, China; Department of Neurology, Xuanwu Hospital of Capital Medical University, Beijing 100053, China; Centre for Medical Genetics and Hunan Key Laboratory of Medical Genetics, School of Life Sciences, Central South University, Changsha, Hunan 410008, China; State Key Laboratory of Membrane Biology, Peking University School of Life Sciences, Beijing 100871, China; PKU-IDG/McGovern Institute for Brain Research, Beijing 100871, China; Peking-Tsinghua Center for Life Sciences, Academy for Advanced Interdisciplinary Studies, Peking University, Beijing 100871, China; In Vivo Neurobiology Group, Neurobiology Laboratory, National Institute of Environmental Health Sciences, Research Triangle Park, NC 27709, U.S.A.; Laboratory of Physiologic Studies, National Institute on Alcohol Abuse and Alcoholism, National Institutes of Health, Bethesda, MD 20892, U.S.A.; Laboratory for Integrative Neuroscience, National Institute on Alcohol Abuse and Alcoholism, National Institutes of Health, Rockville, MD, 20852, USA; Computational Biology Group, Laboratory of Neurogenetics, National Institute on Aging, National Institutes of Health, Bethesda, MD 20892, U.S.A.; Department of Neurology, Union Hospital, Tongji Medical College, Huazhong University of Science and Technology, Wuhan, Hubei 430022, China; Department of Neurosciences, University of South China Medical School, Hengyang, China; Chinese Institute for Brain Research, Beijing 102206, China; National Clinical Research Center for Geriatric Disorders, Xiangya Hospital, Central South University, Changsha, Hunan 410008, China

**Keywords:** Parkinson’s disease, substantia nigra, dopaminergic neurons, endocannabinoid, diacylglycerol lipase β, 2-arachidonoyl-glycerol, motor skill learning, motor control, genetics, pathophysiology

## Abstract

2-arachidonoyl-glycerol (2-AG), the most abundant endocannabinoid (eCB) in the brain, regulates diverse neural functions. However, whether 2-AG deficiency contributes to Parkinson’s disease (PD) and nigral dopaminergic neurons (DANs) dysfunction is unclear. Diacylglycerol lipase α and β (*DAGLA* and *DAGLB*) mediate the biosynthesis of 2-AG. Using homozygosity mapping and whole-exome sequencing, we linked multiple homozygous loss-of-function mutations in *DAGLB* to a form of early-onset autosomal recessive PD. We then used RNA sequencing and fiber photometry with genetically encoded eCB sensors to demonstrate that DAGLB is the main 2-AG synthase in nigral DANs. Genetic knockdown of *Daglb* by CRISPR/Cas9 in mouse nigral DANs substantially reduces 2-AG levels in the *substantia nigra* (SN). The SN 2-AG levels are markedly correlated with the vigor of movement during the acquisition of motor skills, while *Daglb*-deficiency impairs motor learning. Conversely, pharmacological enhancement of 2-AG levels increases nigral DAN activity and dopamine release and improves motor learning. Together, we demonstrate that *DAGLB*-deficiency contributes to the etiopathogenesis of PD, reveal the importance of *DAGLB*-mediated 2-AG biosynthesis in nigral DANs in regulating neural activity and dopamine release, and provide preclinical evidence for the beneficial effects of 2-AG augmentation in PD treatment.

## Introduction

Parkinson’s disease (PD) is clinically manifested with both motor and non-motor symptoms ^1^. A preferential degeneration of nigral dopaminergic neurons (DANs) in the ventral *substantia nigra pars compacta* (SNc) and the resulting impairments of dopamine transmission in basal ganglia are broadly responsible for the motor symptoms, which include bradykinesia, resting tremor, rigidity, and motor skill learning deficits ^2,3^. Both genetic and environmental factors contribute to the etiopathogenesis of PD. The identification of monogenetic mutations responsible for various familial forms of PD provide molecular clues in understanding the pathophysiological mechanisms of the disease ^4^. To date, more than 20 genes have been linked to the familial forms of Parkinsonism, including both autosomal dominant and recessive mutations ^5^. However, a significant proportion of familial PD cases are still genetically unexplained. Identifying those unknown genetic factors may uncover new signaling pathways critical for regulating nigral DAN activity and PD pathogenesis.

The nigral DANs are essential in regulating the vigor of movement ^6^ and motor learning ^7^. The activity of nigral DANs and dopamine release can be dynamically regulated by diverse presynaptic inputs, of which the direct pathway striatal spiny projection neurons (dSPNs) in dorsal striatum provide the major inhibitory inputs ^7-11^. High levels of cannabinoid receptor 1 (CB1) are present in the axon terminals of dSPNs ^12,13^, which may respond to the endocannabinoid (eCB) 2-arachidonoyl-glycerol (2-AG) and anandamide (AEA) released from the nigral DANs. The eCBs act as neuromodulators, retrogradely suppress presynaptic neurotransmitter release through the G protein-coupled CB1 receptors, and regulate a variety of physiological processes, such as motor learning, stress response, and memory ^14-18^. The midbrain DANs can produce and release eCBs from soma and dendrites ^19^. Both diacylglycerol lipase α (DAGLA) and its homologue DAGLB mediate the biosynthesis of 2-AG, the most abundant eCB in the brain ^20^. While DAGLA catalyzes most of the 2-AG production in the brain ^21-23^, the main 2-AG synthase in nigral DANs remains to be determined. Confounding upregulation and downregulation of eCBs and receptors have been observed in the basal ganglia of PD patients and related animal models ^17,24,25^. However, it is unclear whether the altered eCB signaling contributes to the disease or merely reflects compensatory responses. To understand how eCB system regulates dopamine transmission in motor control may provide new insights into the pathogenic mechanisms and treatment of PD.

In supporting a critical involvement of eCB signaling in regulating nigral DAN activity and PD pathogenesis, here we first provided genetic evidence to demonstrate that deficiency in 2-AG synthase *DAGLB* contributes to the etiopathogenesis of PD. We then revealed a previously undescribed, nigral DAN-specific pathogenic mechanism of *DAGLB* dysfunction in motor learning. Finally, we showed that pharmacological augmentation of 2-AG levels may serve as a potential therapeutic treatment for PD.

## Results

### Identification of *DAGLB* mutations in patients with early-onset autosomal recessive PD

Previously, we recruited a large cohort of patients with autosomal recessive PD (ARPD) and sporadic early-onset PD (EOPD) in China and identified pathogenic variants of known PD genes in 65 ARPD families using exon dosage analysis and whole-exome sequencing (WES) ^26^. To discover new causal genetic mutations in the remaining ARPD families, we first studied one consanguineous family (Family 1, AR-003) with two siblings affected by EOPD (**Fig. 1A**). Genome-wide single nucleotide polymorphism (SNP) analysis and homozygosity mapping of the affected individuals revealed five regions of homozygosity shared by the affected sisters (II-3 and II-4) as the candidate causative gene regions (**Supplementary Table S1, Supplementary Fig. S1**). Assuming recessive mode of inheritance, we then analyzed the WES data from those two affected siblings and searched for shared homozygous mutations. Consequently, we identified one homozygous splice-site mutation (c.1821-2A>G) residing in intron 14 of *DAGLB* confirmed by Sanger sequencing and segregated with disease in this family (**Fig. 1A, Supplementary Tables S2 and S3, Supplementary Fig. S2**). The “c.1821-2A>G” mutation was predicted in silico to disrupt the donor splice site of exon 15 and confirmed by reverse-transcription PCR analysis (**Supplementary Fig. S3**). Next, we analyzed the *DAGLB* gene for homozygous or compound heterozygous mutations by mining the WES data from an additional 1,654 unrelated PD probands, including 156 ARPD and 1,498 sporadic EOPD cases without known PD-related genetic mutations. Accordingly, we identified one homozygous missense mutation [c.1088A>G (p.D336G)] in Family 2 (AR-005) and one homozygous frameshift mutation [c.469dupC (p.L158Sfs*17)] in Family 4 (**Fig. 1A, Supplementary Tables S2 and S3, Supplementary Fig. S2**). Genome-wide SNP array genotyping also showed homozygosity present in the affected cases from Families 2 and 4, which include the *DAGLB* gene (**Supplementary Fig. S1**). Finally, we performed copy number variation analysis of the WES data and identified one more homozygous deletion (g.ch7:6,486,383-6,489,136del), which contains the entire exon 1 and 5’-untranslated region of *DAGLB* gene in another family with two affected siblings (Family 3, AR-075) and was validated through Oxford Nanopore long-read sequencing and Sanger sequencing (**Fig. 1A, Supplementary Figs. S4 and S5**). These *DAGLB* mutations are absent from or present in the heterozygous state in available unaffected family members and healthy control subjects, strongly supporting the pathogenicity of homozygous *DAGLB* mutations in EOPD. Those six affected individuals had early disease onset (≤ 40 years old) and displayed typical parkinsonism and good levodopa response (**Supplementary Table S4, Supplementary Clinical description**). However, compared to patients with PD-related *PRKN, PINK1* or *DJ-1* recessive mutations, *DAGLB* mutation-affected individuals showed more severe motor manifestations and more non-motor signs, such as depression. Positron emission tomography (PET) revealed impaired dopamine transmission in the striatum (**Supplementary Fig. S6**). Together, we identified four different homozygous *DAGLB* mutations in six affected EOPD individuals. *DAGLB* is the fourth most frequent ARPD gene after *PRKN, PINK1* and *PLA2G6* in our large Chinese APRD cohort ^26^.

**Fig. 1.**
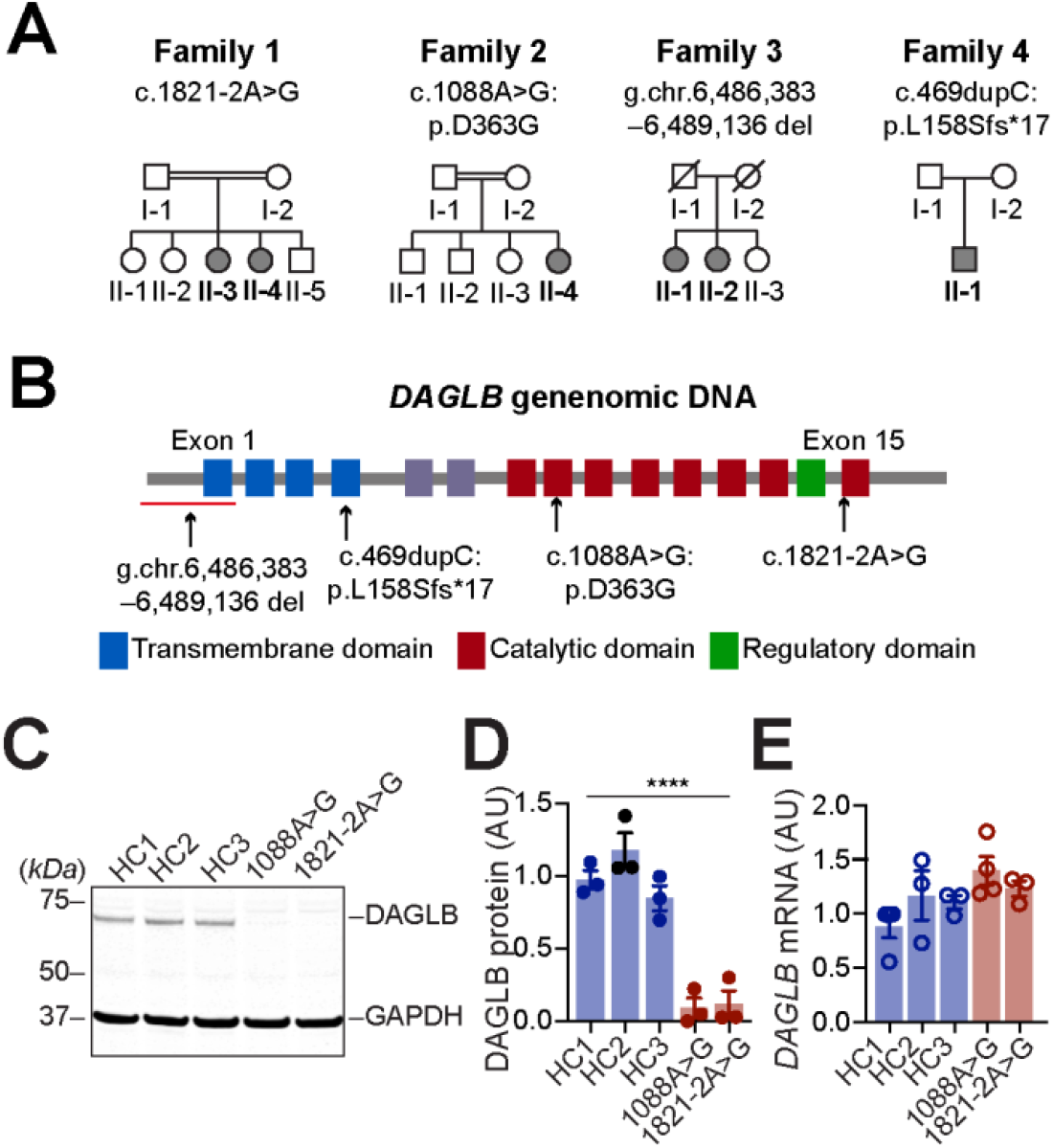
Identification of homozygous *DAGLB* mutations in affected families. **(A)** Pedigrees are shown for the four affected families. A double bar represents parental consanguinity. Slash indicates deceased individuals. Males are represented by squares, females by circles and affected individuals by shading. **(B)** Schematic view of *DAGLB* gene structure and encoded protein domains. The four transmembrane segments are shown in blue, and the catalytic domain in maroon. Within the catalytic domain a regulatory loop is colored in green. Variant sites are indicated by arrows. **(C-E)** Representative western blot **(C)** and quantification of *DAGLB* protein **(D)** and mRNA **(E)** levels in fibroblasts derived from Family 1 II-3, Family 2 II-4 and three age-matched HCs. Data were normalized to glyceraldehyde 3-phosphate dehydrogenase (GAPDH) protein or mRNA levels as appropriate and represent mean ± SEM. ^****^*p* < 0.0001.

### The PD-related mutations disrupt the formation and stability of DAGLB proteins

*DAGLB* encodes a 672-amino acid protein containing four transmembrane domains and one catalytic domain that mediates the biosynthesis of 2-AG ^23^. The “g.ch7:6,486,383-6,489136del” and “c.469dupC” mutations apparently disrupt the translation of the catalytic domain (**Fig. 1B**), resulting in loss-of-function of DAGLB. By contrast, the “c.1821-2A>G” mutation truncates part of the catalytic domain, while the “c.1088A>G” mutation replaces a conserved aspartate (D) residue with glycine (G) in the catalytic domain (**Fig. 1B, Supplementary Fig. S7**). To investigate how the two missense mutations affect the expression of DAGLB protein, we examined the expression of DAGLB protein and *DAGLB* mRNA in primary fibroblasts derived from patients carrying the mutations. Compared to the healthy controls (HC), DAGLB protein was barely detectable by western blot in the patients’ samples [1way ANOVA, F(4, 10)=33.1, p<0.0001] (**Fig. 1C, D**). In contrast, the *DAGLB* mRNA expression is comparable between the patients and control samples [1way ANOVA, F(4, 10)=2.3, p=0.12] (**Fig. 1E**), suggesting that the mutations affect the stability of DAGLB protein. Indeed, treatment with proteasome inhibitor MG132 increased the levels of DAGLB protein in both control and patients’ samples (**Supplementary Fig. S8A, B**). Additionally, the mutations did not affect the expression of DAGLA protein in patients’ samples (**Supplementary Fig. S8C, D**). Therefore, all four PD-related mutations disrupt the formation or stability of DAGLB protein, suggesting that the impairment of *DAGLB*-mediated 2-AG signaling may contribute to the etiopathogenesis of PD.

### *DAGLB* is the main 2-AG synthase expressed in nigral DANs

While our human genetic studies linked deficiency in *DAGLB* to PD (**Fig. 1**), *DAGLA* is the main 2-AG synthesis in the CNS and account for 80% production of 2-AG in the mouse brains ^21,22^. How does the *DAGLB*-deficiency contribute to PD and nigral DAN dysfunction? Interestingly, a previous whole genome RNA-sequencing study ^27^ revealed 10-fold more abundance of *DAGLB* than *DAGLA* mRNA in laser capture microdissection (LCM)-isolated human nigral DANs (**Fig. 2A**, unpaired *t* test, p<0.0001, plotted from GSE76514). We then performed RNA-sequencing of LCM-isolated mouse nigral DANs and found that the expression of *Daglb* mRNA is 2-fold higher than *Dagla* (unpaired *t* test, p<0.0001) (**Fig. 2B**). By contrast, *Dagla* mRNA was more enriched in striatal neurons than *Daglb* (unpaired *t* test, p<0.0001) (**Fig. 2C**). To determine the cellular location of DAGLB protein in nigral DANs, we tested the commercially available DAGLB antibodies. Unfortunately, none of them stained midbrain DANs. However, RNAscope^®^ *in situ* hybridization demonstrated the co-localization of *Daglb* and *Dagla* mRNA with the dopamine synthase *tyrosine hydroxylase* (*Th*) in mouse nigral DANs (**Fig. 2D, Supplementary Fig. S9**), Therefore, while *DAGLA* is highly expressed by most neurons in the brain ^21-23^, *DAGLB* is the main 2-AG synthase expressed by nigral DANs, suggesting a nigral DAN-specific mechanism of *DAGLB*-deficiency in PD.

**Fig. 2.**
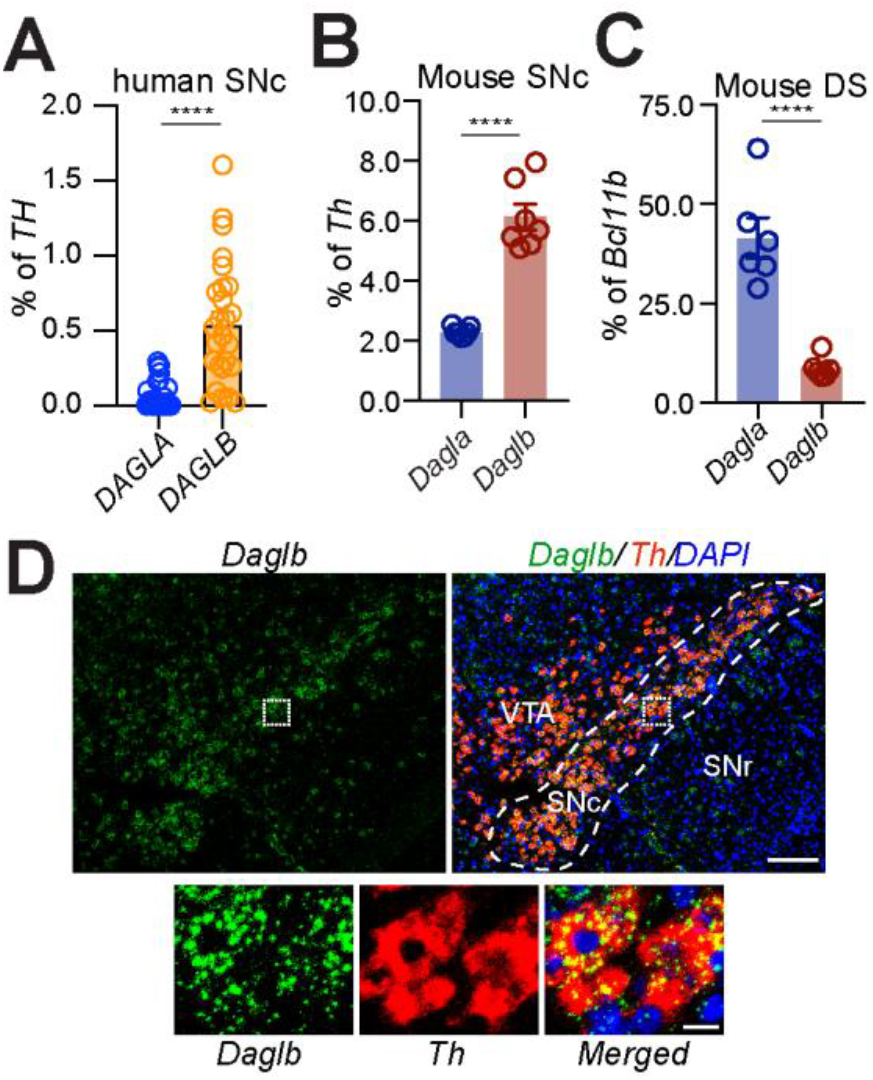
DAGLB is the main 2-AG synthase expressed by the nigral DANs. (**A**) Quantification of *DAGLA* and *DAGLB* mRNA expression by RNA-sequencing of LCM-isolated human nigral DAN samples (n=26) ^27^. **(B)** Quantification of *Dagla* and *Daglb* mRNA expression by RNA-sequencing of LCM-isolated nigral DANs from adult mouse brains (n = 7). **(C)** *Dagla* and *Daglb* mRNA expression in the mouse dorsal striatum (DS, n=6). The expression of *Dagla* and *Daglb* mRNAs was normalized by the expression of *Th* mRNAs in the nigral DANs and *Bcl11b* mRNAs in the SPNs. Data were presented as mean ± SEM. ^****^*p*<0.0001. **(D)** RNAscope^®^ *in situ* hybridization of *Daglb* and *Th* in mouse midbrain sections. Sections were counterstained with DAPI. Dashed line outlines the SNc region. Right panels highlight the boxed areas in the left panels. SNc: substantia nigra pars compacta. SNr: substantia nigra pars reticulata. VTA: ventral tegmental area. Scale bars: 100 μm (left) and 20 μm (right).

### *Daglb*-knockdown in nigral DANs reduces 2-AG levels in the SN

To examine the functional significance of DAGLB, we decided to selectively knockdown (KD) *Daglb* or *Dagla* in nigral DANs using an adeno-associated virus (AAV)-based CRISPR/SaCas9 genome editing system (AAV-CMV-DIO-SaCas9-U6-sgRNA) ^28^. The control (Ctrl) saCas9 empty (referred to as the AAV-Ctrl) and saCas9 with the *Daglb* or *Dagla*-guided sgRNA (referred to as the AAV-*Daglb* KD and AAV-*Dagla* KD) are expressed in a Cre-dependent manner (**Fig. 3A**). Co-transfection of AAV-Cre and AAV-*Daglb* KD vectors led to substantial reduction of DAGLB protein levels but not DAGLA in primary cultured mouse cortical and hippocampal neurons, while co-transfection of AAV-Cre and AAV-*Dagla* KD vectors specifically suppressed the expression of DAGLA protein (**Fig. 3B-D**). Therefore, we developed *Daglb*- and *Dagla*-specific genetic KD AAV vectors to selectively manipulate the levels of *Daglb* and *Dagla* expression in a cell-type dependent manner.

**Fig. 3.**
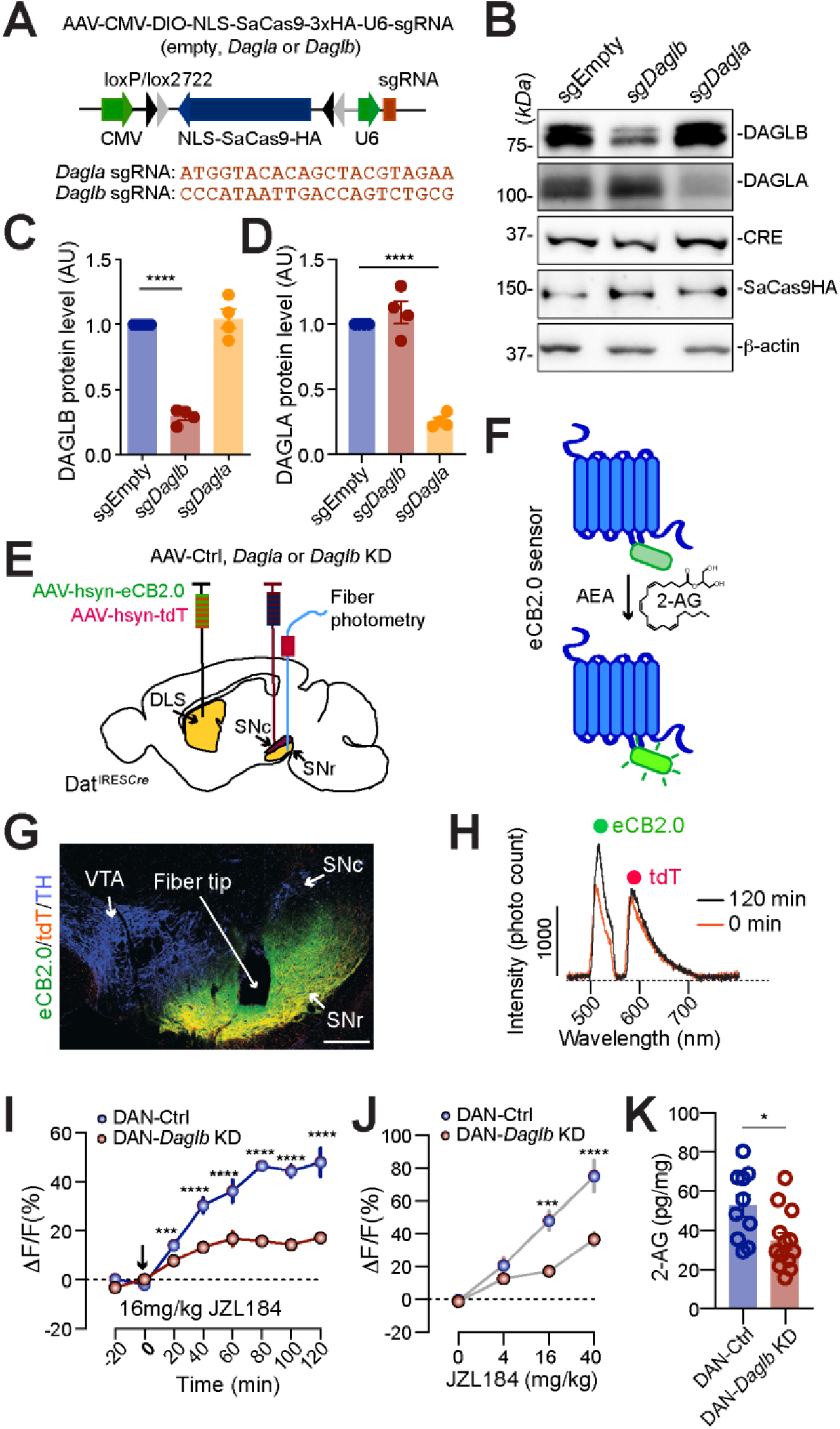
DAGLB mediates the main 2-AG synthesis in the nigral DANs. **(A)** Diagram of AAV-mediated CRISPR/saCas9 gene targeting vector and the sequence of *Dagla* and *Daglb* sgRNAs. **(B)** Western blot of DAGLB in cultured cortical and hippocampal neurons transfected with control, *Dagla*, or *Daglb* KD AAV vectors in combination with Cre-expressing AAV vectors. Actin was used as a loading control. **(C, D)** Bar graph quantifies DAGLB (**C**) and DAGLA (**D**) protein levels in four independent experiments. Data were presented as mean ± SEM. Tukey’s multiple comparison test, ^****^*p*<0.0001. **(E)** Schematic illustrates fiber photometry imaging of 2-AG signals in the SN of DAN-control and DAN-*Daglb* KD mice. DLS: dorsolateral striatum. **(F)** Cartoon of eCB sensor eCB2.0. **(G)** Co-staining of eCB2.0, tdT and TH in the midbrain sections of DAN-*Daglb* KD mice. Scale bar: 100 μm. **(H)** Sample photon counts of eCB2.0 (wavelength: 500-540nm) and tdT (wavelength: 575-650nm) emission immediately before and 120 min after JZL184 (16mg/kg) administration. **(I)** Time course of eCB2.0 signals in the SN of DAN-control [n=4, 2Male(M)/2Female(F)] and DAN-*Daglb* KD (n=4, 2M/2F) mice before and after JZL184 (16mg/kg) treatment. Data were presented as mean ± SEM. Sidak’s multiple comparison test. ^***^*p*=0.0001. ^****^*p*<0.0001. **(J)** Dose response of eCB2.0 signals 120 min after JZL administration at 0 (vehicle only), 4, 16, and 40mg/kg. n=4 mice per group. Data were presented as mean ± SEM. Sidak’s multiple comparison test. ^***^*p*=0.0003. ^****^*p*<0.0001. (**K**) LC-MS/MS quantification of 2-AG in the SNc of DAN-control (n=10) and DAN-*Daglb* KD (n=14) mice. Data were presented as mean ± SEM. Unpaired t test, ^*^*p*=0.01.

To measure 2-AG release in the SN *in vivo*, we stereotactically injected the AAVs carrying a genetically encoded eCB sensor named eCB2.0 ^29,30^ in the dorsal striatum (**Fig. 3E, F**). A custom-built fiber photometry system ^31^ was employed to capture the eCB2.0 signals in the dSPN-projecting *substantia nigra pars reticulata* (SNr) area (**Fig. 3E**), where the axons of dSPNs and dendrites of DANs form synaptic connections ^10,13^. The infusion of control (Ctrl)-, *Daglb* KD- or *Dagla* KD-AAVs in the SNc of DAT^IRES*Cre*^ mice leads to selective expression of either saCas9 empty (referred to as DAN-Ctrl) or saCas9 with the *Daglb*- or *Dagla*-sgRNA (referred to as DAN-*Daglb* KD or *Dagla* KD) in the DANs (**Fig. 3E**). Overall, around 75% of DANs in the SNc were infected with AAV-Ctrl or AAV-*Daglb* KD, while no apparent loss of nigral DANs was observed in the DAN-*Daglb* KD mice 12 months after AAV injection (**Supplementary Fig. S10A-D**).

The eCB2.0 sensor was constructed based on CB1 receptors, of which the third intracellular loop is replaced with circularly permuted green fluorescent protein (cpGFP) and the binding of 2-AG and AEA enhances the emission intensity of cpGFP ^29^ (**Fig. 3F**). Like the native CB1 receptor, eCB2.0 sensors were transported to the dSPN axon terminals in the SN (**Fig. 3G**). The AAVs carrying red fluorescent protein tdTomato (tdT) were co-injected with AAV-eCB2.0 as a reference for adjusting any motion artifacts during the imaging process ^31^. Two distinct emission peaks corresponding to the eCB2.0 and tdT signals were detected in the SNr immediately before and 120 min after the administration of JZL184 (16mg/kg), a selective monoacylglycerol lipase inhibitor ^32^ that blocks the degradation of 2-AG (**Fig. 3H**). The JZL184-induced enhancement of eCB2.0 signals was substantially diminished in the DAN-*Daglb* KD mice compared to the controls in both time- and dose-dependent manners [time: 2way ANOVA, F(7,42)=60.3, p<0.0001; dose: 2way ANOVA, F(3,18)=66.7, p<0.0001] (**Fig. 3I, J**). Furthermore, liquid chromatography-tandem mass spectrometry (LC-MS/MS) also revealed a marked reduction of 2-AG levels in the SNc of DAN-*Daglb* KD mice (unpaired *t* test, p=0.01) (**Fig. 3K**). By contrast, genetic deletion of *Dagla* in the SN DANs did not affect the JZL184-induced enhancement of eCB2.0 signals in the SNr (**Supplementary Fig. S11)**. Together, these results demonstrate that DAGLB is the dominant 2-AG synthase that catalyzes the 2-AG production in nigral DANs.

### The nigral 2-AG levels correlate with motor performance during motor skill learning

Our recent study demonstrates that ablation of nigral DANs in mouse models only modestly reduced the walking speed, but completely abolished the improvement of motor performance in rotarod motor skill learning test ^7,33^, revealing a critical involvement of nigral DAN activity in motor skill learning. We thereby examined the 2-AG signals in the SNr by simultaneously conducting fiber photometry live recording and the rotarod training, 10 trials per session on each day for six consecutive days **(Fig. 4A)**. Each trial started with a low constant rotating speed at 4 rpm for 30 sec before linear acceleration from 4 to 40 rpm in 5 min ^34^. The 2-AG signals gradually increased along with the progression of 10 trials during each training session (**Fig. 4A, B, Supplementary Fig. S12**). In addition, more robust daily enhancement of 2-AG levels was recorded on the first four days’ training compared to the last two days’ [1way ANOVA, F(5,42)=12.2, p<0.0001] (**Fig. 4C**). The first four days’ training is generally regarded as the acquisition phase of motor learning, while the last two days are regarded as the maintenance or retention phase ^35,36^. As expected, rotarod performance was also greatly improved during the acquisition phase, but not in the retention phase **(Fig. 4D)**. Indeed, further correlational analyses reveal stronger positive correlations between the 2-AG signal enhancement and rotarod performance in the acquisition phase compared to the retention phase [1way ANOVA, F(5,42)=16.9, p<0.0001] (**Fig. 4E**). These results suggest that the SN 2-AG signaling is particularly engaged with the vigor of movement during the acquisition phase of motor skill learning.

**Fig. 4.**
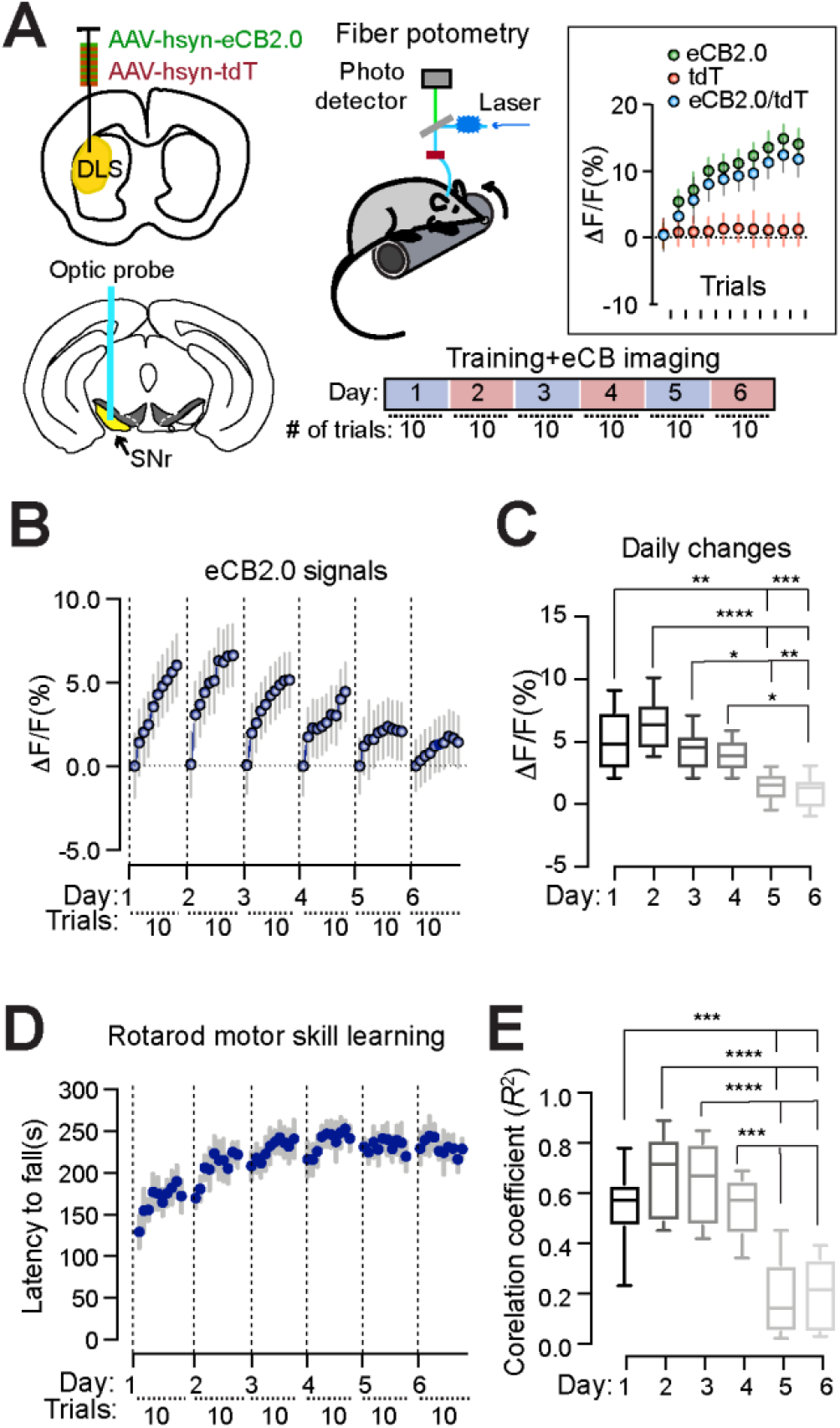
DAGLB-mediated 2-AG biosynthesis in the nigral DANs is involved in motor skill learning. **(A)** Schematic diagram of eCB2.0 fiber photometry recoding while mice undergo rotarod motor skill training. Inset shows the average eCB2.0 (green), tdT (red) and normalized eCB2.0 (blue) signals during each trial from a mouse on day 2 of the 6-day training paradigm. Data were presented as mean ± SD. **(B)** Normalized eCB2.0 signal intensity during the motor learning. n=8 (4M/4F). Data were presented as mean ± SD. **(C)** Box and whiskers plot (min to max) of maximal daily increase of eCB2.0 signal intensity. n=8 (4M/4F). Tukey’s multiple comparisons test. ^*^*p*=0.02, ^**^*p*=0.001, ^***^*p*=0.0003, ^****^*p*<0.0001. **(D)** Performance of rotarod motor skill training. n=8 (4M/4F). Data were presented as mean ± SEM. **(E)** Box and whiskers plot (min to max) of correlation coefficient of eCB2.0 signal intensity and rotarod performance. n=8 (4M/4F). Tukey’s multiple comparisons test. ^***^*p*=0.0002 to 0.0006, ^****^*p*<0.0001.

### *Daglb*-knockdown in the nigral DANs compromises the dynamic 2-AG release during the early phase of motor skill learning and impairs the overall motor performance

To further examine the contribution of *Daglb*-mediated 2-AG biosynthesis in the nigral DAN-dependent motor skill learning, we selectively knock-downed the expression of *Daglb* in nigral DANs of 3-month-old DAT^IRESCre^ mice by AAV vectors and compared the SN 2-AG levels in DAN-Ctrl and DAN-*Daglb* KD mice during rotarod tests (**Fig. 5A**). The increase of eCB2.0 signals was less robust in the SN of DAN-*Daglb* KD mice compared to the DAN-Ctrl ones, especially in the acquisition phase of motor learning [2way ANOVA, genotype: F(1, 8)=13. 68, p=0.006] (**Fig. 5A**). Correlatively, the DAN-*Daglb* KD mice displayed marked impairments in the overall performance of rotarod motor learning tests compared to the control mice [2way ANOVA, genotype: F(1,30)=9.3, p=0.0047] (**Fig. 5B**). By contrast, there were no apparent alterations of spontaneous locomotor activity nor gait properties of DAN-*Daglb* KD mice compared to the controls (**Supplementary Fig. S13**). Together, these results demonstrate that the DAGLB-mediated 2-AG biosynthesis in the nigral DANs is actively engaged in regulating the functional role of nigral DANs in motor skill learning.

**Fig. 5.**
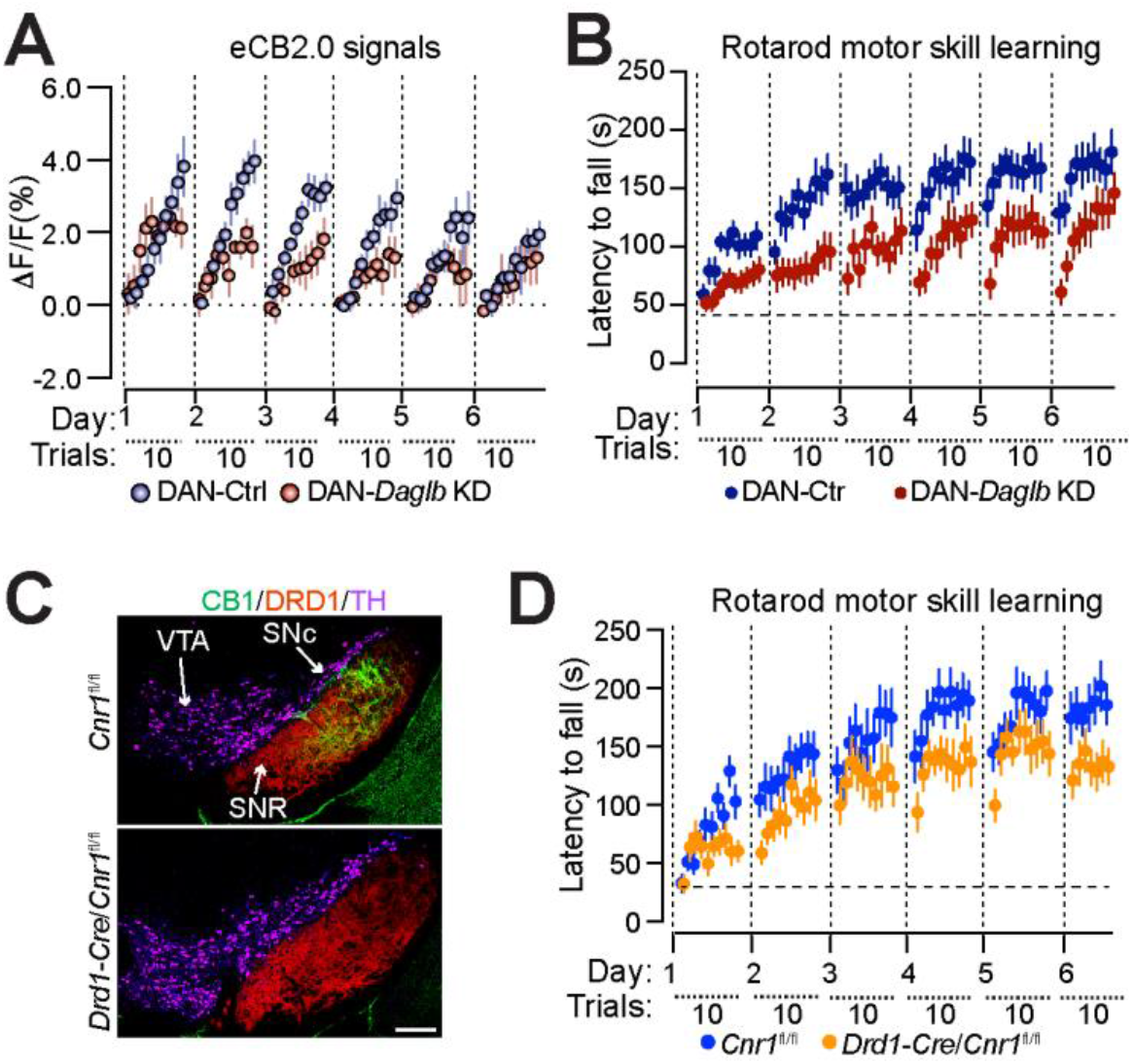
*DAGLB*-deficiency in nigral DANs compromises the early dynamic release of 2-AG and impairs rotarod motor skill learning. **(A)** Normalized eCB2.0 signal intensity over the course of motor learning of DAN-control and DAN-*Daglb* KD (n=5M per genotype) mice. Data were presented as mean ± SEM. **(B)** Rotarod motor skill training of DAN-control (n=16, 8M/8F) and DAN-*Daglb* KD (n=16, 8M/8F) mice. Data were presented as mean ± SEM. **(C)** Co-staining of CB1 (green), DRD1 (red) and TH (purple) in the midbrain sections of homozygous floxed CB1 (*Cnr1*^fl/fl^) and Drd1-Cre/*Cnr1*^fl/fl^ mice. Scale bar: 100 μm. **(D)** Rotarod motor skill training of *Cnr1*^fl/fl^ (n=14, 7M/7F) and Drd1-Cre/*Cnr1*^fl/fl^ (n=12, 6M/6F) mice. Data were presented as mean ± SEM.

We next examined whether CB1 receptors in the axon terminals of dSPNs mediate the retrograde 2-AG signaling during motor skill learning. To selectively delete the CB1 receptor-encoding *Cnr1* gene in the dSPNs, we crossbred a line of *Cnr1*-floxed (*Cnr1*^fl/fl^) mice ^38^ with dopamine receptor 1-Cre (*Drd1*-Cre) mice. Accordingly, the expression of CB1 receptors was completely abolished in the SNr of *Drd1*-Cre/ *Cnr1*^fl/fl^ bigenic mice (**Fig. 5C**). Similarly to the DAN-*Daglb* KD mice, the *Drd1*-Cre/ *Cnr1*^fl/fl^ mice also displayed impairments in rotarod motor skill learning compared to the littermate controls [2way ANOVA, genotype: F(1, 24)=6.301, p=0.0192] (**Fig. 5D**). Therefore, the nigral DAN-derived 2-AG likely regulates motor learning through modulating the presynaptic inputs from dSPNs.

### *Daglb* germline knockout mice do not developed any overt motor behavioral and neuropathological abnormalities

Like *DAGLB*, the loss-of-function mutations in *PARKIN, DJ-1*, and *PINK1* also contribute to the etiopathogenesis of PD ^5^. However, the *Parkin, Dj-1*, and *Pink1* germline knockout (KO) mice did not develop any overt PD-related behavioral and neuropathological abnormalities ^37^. Similarly, we did not observe any apparent alterations of locomotor activity in *Daglb* germline KO mice at 4, 8, 12, and 20 months of age (**Supplementary Fig. S14A-C**). Since the rotarod motor learning test provides a more sensitive behavioral paradigm to detect the dysfunction of nigral DANs ^7^, we examined the rotarod performance of *Daglb* germline KO mice at 4 and 20 months of age. The 4-month-old *Daglb* KO mice displayed modest but statistically insignificant improvement of motor learning [2way ANOVA, genotype: F(1, 19)=2.9, p=0.103] (**Supplementary Fig. S14D**), while the 20-month-old *Daglb* KO mice showed similar performance compared to the littermate controls [2way ANOVA, genotype: F(1, 19)=0.02, p=0.899] (**Supplementary Fig. S14E**). Additionally, no apparent loss of TH-positive nigral DANs was found in the 20-month-old *Daglb* KO mice [Unpaired *t* test, p=0.9989] (**Supplementary Fig. S14F**). Therefore, the CRISPR/SaCas9-mediated acute knockdown of *Daglb* in the nigral DANs of adult mice provide a more sensitive experimental paradigm to evaluate the contribution of DAGLB-dependent 2-AG signaling in nigral DANs during motor learning.

### 2-AG signaling potentiates nigral DAN activity and somatodendritic dopamine release

Since 2-AG from nigral DANs acts on the presynaptic CB1 receptors to suppress the release of inhibitory neurotransmitter GABA from dSPN axon terminals ^19^, we suspected that the JZL184-induced 2-AG upregulation in the SN (**Fig. 3I, J**) may disinhibit the presynaptic inhibitory inputs from dSPNs and lead to enhanced DAN activity and somatodendritic dopamine release. To test this hypothesis, we first treated the mice with JZL184, and then used fiber photometry with genetically encoded calcium indicator GCaMP6f ^39^ and dopamine indicator DA2m ^40^ to monitor the DAN activity and somatodendritic dopamine release. To examine the DAN calcium transients, we crossbred DAT^IRESCre^, Ai95 (RCL-GCaMP6f) and Ai9 (RCL-tdT) mice to selectively express GCaMP6f and tdT in the midbrain DANs of DAT^IRESCre^/GCaMP6f/tdT trigenic mice, and then stereotaxically injected AAV-Ctrl or AAV-*Daglb* KD vectors in the SNc of trigenic mice to evaluate the role of DAGLB in regulating DAN activity (**Fig. 6A, B**). The intraperitoneal injection of JZL184 (20 mg/kg) led to substantial increase of DAN activity as indicated with the elevated GCaMP6f signal intensities in the SNc of both DAN-Ctrl [2way ANOVA, treatment: F(1, 4)=36.89, p=0.0037] and DAN-*Daglb* KD [2way ANOVA, treatment: F(1, 4)=36.88, p=0.0037] trigenic mice compared to the vehicle treatment (**Fig. 6C**). However, the JZL184 treatment induced less robust enhancement of neural activity [2way ANOVA, genotype: F(1, 4)=14.58, p=0.0188] in the SNc of DAN-*Daglb* KD mice compared to the DAN-Ctrl mice (**Fig. 6C**).

**Fig. 6.**
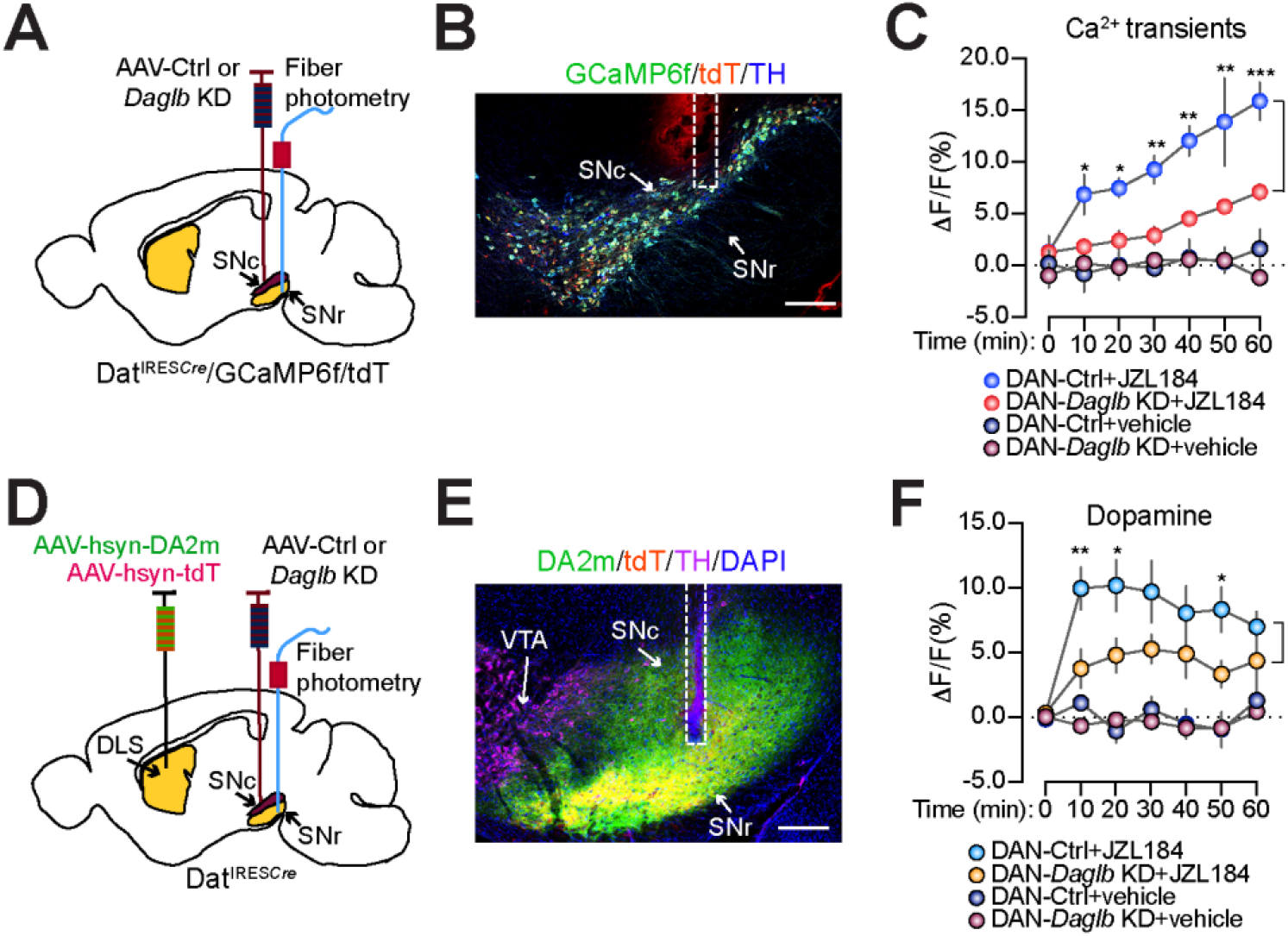
JZL184 treatment promotes DAN activity and dopamine release. **(A)** Schematic illustrates fiber photometry imaging of GCaMP6f and tdT signals in the SNc of DAN-control and DAN-*Daglb* KD Dat^IRESCre^/GCaMP6f/tdT trigenic mice. (**B**) Representative images of GCaMP6f (green), tDT (red) and TH (blue) staining. Scale bar: 200μm. **(C)** Alterations of calcium transients in the SNc of DAN-control and DAN-*Daglb* KD Dat^IRESCre^/GCaMP6f/tdT trigenic mice (n=3M per genotype) treated with vehicle or JZL184 (20mg/kg). Data were presented as mean ± SEM. Multiple *t* test. ^*^*p*<0.05, ^**^*p*<0.01, ^***^*p*<0.001. **(D)** Schematic illustrates fiber photometry imaging of DA2m and tdT signals in the SN of DAN-control and DAN-*Daglb* KD mice. (**E**) Representative images of DA2m (green), tDT (red) and TH (magenta) staining. Scale bar: 200μm. **(F)** Changes of dopamine release in the SN of DAN-control (n=5M) and DAN-*Daglb* KD (n=4M) mice treated with vehicle, JZL184 (20mg/kg). Data were presented as mean ± SEM. Multiple *t* test. ^*^*p*<0.05, ^**^*p*<0.01.

To monitor dopamine release in the SN of DAN-Ctrl and DAN-*Daglb* KD mice, we stereotaxically injected AAV-DAm2 and AAV-tdT vectors in the dorsal striatum, and AAV-Ctrl or AAV-*Daglb* KD vectors in the SNc of DAT^IRESCre^ mice (**Fig. 6D, E**). The same JZL184 treatment also substantially enhanced dopamine release as indicated with the increased DA2m fluorescent signal intensities in the SN of both DAN-Ctrl [2way ANOVA, treatment: F(1, 8)=23.13, p=0.0013] and DAN-*Daglb* KD [2way ANOVA, treatment: F(1, 6)=18.7, p=0.0049] mice compared to the vehicle treatment. While the JZL184-induced dopamine release was not statistically significant between DAN-*Daglb* KD and DAN-Ctrl mice during the entire 60 min period [2way ANOVA, genotype: F(1, 7)=4.612, p=0.0689], multiple comparisons showed markedly less dopamine levels in the SN of DAN-*Daglb* KD mice after the drug treatment in 10, 20, and 50 min (**Fig. 6F**). These data suggest a dynamic interplay between the dopamine and 2-AG signaling in the nigral DANs, in which the DAGLB-mediated 2-AG biosynthesis in nigral DANs promotes the DAN activity and somatodendritic dopamine release.

### Inhibition of 2-AG degradation rescues the motor impairments of *Daglb*-deficient mice

Since the JZL184 treatment enhanced DAN activity and dopamine release in both the control and DAN-*Daglb* KD mice (**Fig. 6C, F**), and the activity of nigral DANs are essential for motor skill learning ^7,33^, we then treated the mice with JZL184 or vehicle 1 hour before each day’s rotarod motor training sessions. The JZL184 treatment (20 mg/kg) markedly improved the motor learning in both DAN-*Daglb* KD mice [2way ANOVA, treatment: F(1, 21)=59.9, p<0.0001] and DAN-Ctrl mice [2way ANOVA, treatment: F(1, 15)=18.3, p<0.0001] compared with the vehicle-treated ones (**Fig. 7**). Moreover, the administration of JZL184 completely rescued the motor learning deficits of DAN-*Daglb* KD mice and made those mice performed even better than the vehicle-treated DAN-Ctrl mice [2way ANOVA, genotype: F(1, 17)=6.645, p=0.0196] (**Fig. 7**). Therefore, the blockage of 2-AG degradation by JZL184 is sufficient to restore the local 2-AG levels required for the rotarod motor skill learning in DAN-*Daglb* KD mice.

**Fig. 7.**
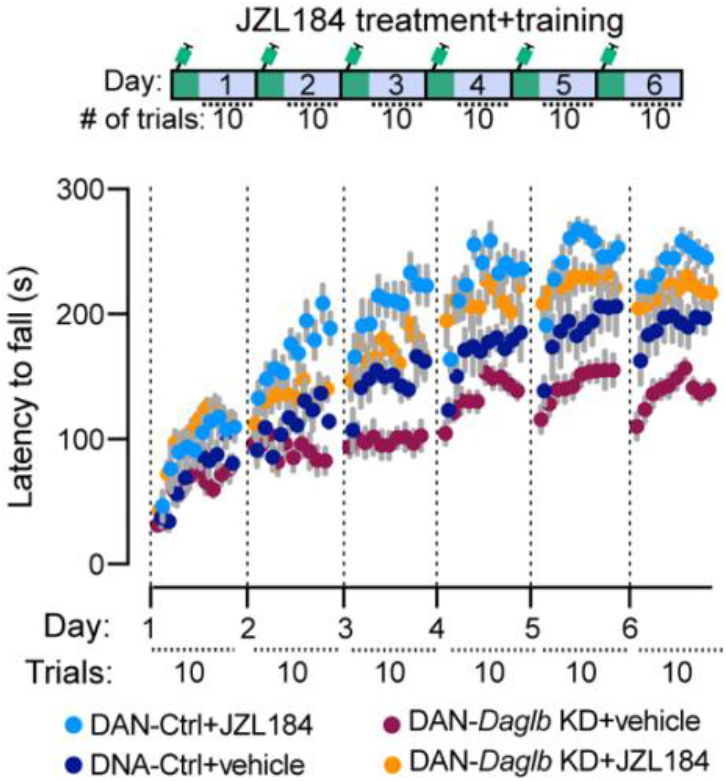
JZL184 treatment rescues the motor skill learning impairments of DAN-*Daglb* KD mice. Rotarod motor skill learning of DAN-control (n_vehicle_=7, 4M/3F; n_JZL184_=10, 5M/5F) and DAN-*Daglb* KD (n_vehicle_=11, 6M/5F; n_JZL184_=12, 6M/6F) mice treated with vehicle or JZL184 (20mg/kg).

## Discussion

In the present work we identified four novel PD-causal loss-of-function mutations in *DAGLB* and showed that DAGLB is the dominant 2-AG synthase in nigral DANs. In supporting the physiological importance of *DAGLB*-mediated 2-AG biosynthesis in nigral DAN-dependent motor functions, we found that genetic knockdown of *Daglb* in the mouse nigral DANs led to reduced nigral 2-AG levels and impaired rotarod motor skill learning, whereas pharmacological inhibition of 2-AG degradation increased nigral 2-AG levels, promoted DAN activity and dopamine release, and rescued the motor deficits. Therefore, we reveal a previously undescribed, DAN-specific pathophysiological mechanism of *DAGLB* dysfunction in PD pathogenesis and provide the rationale and additional preclinical evidence for the beneficial effects of eCB supplementation in PD treatment ^41^.

High levels of eCBs were detected in the cerebrospinal fluid of untreated PD patients ^42^. Increased eCB levels in the globus pallidus are associated with reduced movement in a PD animal model^43^. However, previous studies in PD patients and related animal models mostly focused on the alterations of eCB signaling after severe nigral DAN loss or lengthy levodopa administration ^25^. The results are thereby more likely to reflect the compensatory responses to the disease. It was unclear, however, whether the changes of eCB signaling contribute to the etiopathogenesis of the disease. Our human genetics study provides unequivocal genetic evidence and for the first time demonstrates that like dopamine deficiency, the impairment of 2-AG signaling also contributes to the pathogenesis of PD. *DAGLB* is a gene duplication of *DAGLA*^23^. Although DAGLA is the dominant 2-AG synthase in most neurons and accounts for 80% of 2-AG production in the CNS ^21,22^, our gene expression and functional assays demonstrate that DAGLB mediates the major 2-AG biosynthesis in nigral DANs. The predominant presence of *DAGLB* in nigral DANs may explain why the loss-of-function mutations in *DAGLB* leads to DAN dysfunction and PD. On the other hand, the elevation of 2-AG levels in the other brain regions as observed in the PD patients ^42^ likely represents a compensatory response to the loss of *DAGLB*-mediated 2-AG production in the nigral DANs due to PD-related dopaminergic neurodegeneration.

It might not be totally surprising that *Daglb, Parkin, Dj-1*, and *Pink1* germline KO mice all failed to develop any PD-like behavioral and pathological phenotypes. Longer lifespan and other genetic and physiological characteristics may render human neurons more susceptible to the disease-related mutations ^44^. To overcome the futility in modeling the PD-related recessive mutations with germline KO mice, we applied CRISPR/saCas9-mediate knockdown of *Daglb* selectively in the nigral DANs of adult mice to avoid any potential compensatory interference during development. We also subjected the DAN-*Daglb* KD mice to the nigral DAN-dependent rotarod motor skill learning test to examine any DAN dysfunction. Finally, we employed fiber photometry live recording technique to monitor the 2-AG release in behaving mice in correlation with the motor performance. Together, we offer a new experimental scheme to study the pathophysiological mechanism of PD-related genetic mutations in mouse models, and reveal a new nigral DAN-specific pathogenic mechanism of *Daglb*-deficiency in PD. Since the overall efficiency of CRISPR/saCas9-mediated *Daglb* knockdown is about 70-80% in the current study, it is likely that the residual DAGLB activity in nigral DANs contributes to the increase of 2-AG levels during rotarod motor training and after JZL184 administration. A line of *Daglb* conditional KO mice that selectively delete *Daglb* in the adult nigral DANs would be useful to reveal potentially more severe behavioral and neurochemical phenotypes. In addition, DAGLA, although a minor 2-AG synthase in nigral DANs, may also contribute to the residual 2-AG production in *Daglb*-deficient DANs. Genetic deletion of both *Dagla* and *Daglb* in nigral DANs may provide the means to critically evaluate the pathophysiological role of 2-AG in nigral DAN-dependent motor behaviors.

DAGLA protein is enriched in dendritic spines ^21,22^; however, the subcellular localization of DAGLB protein remains unclear due to a lack of specific antibodies for tissue staining. Considering that the CB1-positive axon fibers form close contact with the dendrites and cell bodies of ventral nigral DANs ^12^ and 2-AG acts within a short range (∼10 μm) from the release sites ^29,30^, it is reasonable to assume that DAGLB is distributed in the soma and dendrites of DANs for local 2-AG production and release. The strenuous rotarod training paradigm appears to promote the somatodendritic release of 2-AG from the nigral DANs more pronouncedly during the early phase of motor learning. The elevated 2-AG likely acts on the presynaptic CB1 receptors to suppress the release of the inhibitory neurotransmitter GABA from the dSPN axon terminals ^19^, resulting in enhanced DAN firing and dopamine release critical for the motor performance and learning process. By contrast, the suppression of *Daglb* expression in nigral DANs dampened the dynamic enhancement of 2-AG release especially during the acquisition of motor skills and compromised the motor performance. Consistently, a previous study also demonstrated that the administration of eCB agonist Δ9-tetrahydrocannabinol increases the DAN firing rate, dopamine synthesis, and dopamine release in dopaminergic axon terminals in striatum ^45^, while the CB1 receptor agonist WIN55,212-2 induces dose-dependent increases in firing rate and burst firing in nigral DANs ^46^. On the other hand, genetic deletion of CB1 receptors in dSPNs completely abolished the expression of CB1 receptors in SNr and induced similar motor learning impairments as the DAN-*Daglb* KD mice. Therefore, the *DAGLB*-mediated 2-AG production in nigral DANs may enhance the nigral DAN activity and somatodendritic dopamine release and facilitate the motor learning and control through attenuating the inhibitory inputs from dSPNs. Since 2-AG works locally near the production and release sites, we only examined the interplay between 2-AG and dopamine signaling in the SN regions. Future experiments will be needed to investigate whether the change of 2-AG release in the soma and dendrites of nigral DANs affects the dopamine release in DAN axon terminals at dorsal striatum.

Although the rotarod motor training paradigm promotes the on-demand 2-AG production by DAGLB, how the DAGLB activity is regulated remains to be determined. Striosome dSPN axons are intermingled with the dendrites of aldehyde dehydrogenase 1a1 (ALDH1A1)-positive DANs perpendicularly protruding in the SNr and form this so-called striosome-dendron bouquet structures ^10,11,13^. The ALDH1A1-positive nigral DANs display distinct rebound activity in response to the inhibitory inputs from dSPNs ^10,11^, which then trigger large dendritic Ca^2+^ transients likely through T-type Ca^2+^ channels ^11,47^. Future studies will be needed to selectively knockout *Daglb* in the ALDH1A1-positive DANs and evaluate whether the intracellular Ca^2+^ elevation in dendrites is required to induce on-site 2-AG production and release, which in turn retrogradely suppress the GABAergic inhibition from dSPNs and further accelerate the rebound activity of nigral DANs. Besides the transsynaptic action, 2-AG can also function cell-autonomously within the DANs through promoting both the pace-maker activity and evoked burst firing ^48^, suggesting that *DAGLB*-deficiency in nigral DANs could also lead to reduced DAN activity and associated motor impairments. Therefore, the nigral DANs can produce and release both dopamine and 2-AG, while 2-AG may further boost the dopamine release and neural activity in response to increasing demand.

The present study focused on the neuronal function of 2-AG; however, 2-AG is also implicated in inflammation ^49^. Myeloid synthesis of 2-AG appears to promote vascular inflammation and atherogenesis ^50^. *Daglb* inactivation in mouse peritoneal macrophages attenuates lipopolysaccharide-induced release of proinflammatory cytokine tumor necrosis factor-α ^51^. Since the inhibition of DAGLB activity works against inflammatory responses ^51^, we reason that the *DAGLB*-deficiency is less likely to directly induce the harmful neuroinflammation implicated in the pathogenesis of PD. Nonetheless, future study will be needed to further elucidate the role of *DAGLB* in microglia or other non-neuronal cells in PD.

In conclusion, our study supports a critical involvement of *DAGLB*-mediated 2-AG biosynthesis in regulating the normal physiological function of nigral DANs, which may help to explain how *DAGLB*-deficiency contributes to PD-related motor symptoms. To boost the production of 2-AG may thereby serve as a potential mechanistic-based therapeutic intervention in PD treatment. Indeed, an exploratory clinical trial of eCB-like cannabidiol seems to improve the mobility and mental states of PD patients ^41^.

## Materials and Methods

### Study participants

Participants were recruited at Xiangya Hospital, Central South University between October 2006 and January 2019 and other hospitals of Parkinson’s Disease and Movement Disorders Multicenter Database and Collaborative Network in China (PD-MDCNC, http://pd-mdcnc.com:3111/) established by our group. These participants include 156 cases with ARPD, 1,498 cases of sporadic EOPD, and 1,758 matched healthy control subjects ^26^. All individuals were subjected to the standard clinical neurological examination. PD was diagnosed according to the UK Parkinson’s disease Society Brain Bank clinical diagnostic criteria ^52^ or Movement Disorders Society (MDS) clinical diagnostic criteria for Parkinson’s disease ^53^ by at least two neurologists. The healthy subjects did not have any nervous system or psychiatric diseases. Human blood samples and fibroblasts were obtained after subjects provided written informed consent. All investigations were conducted according to the Declaration of Helsinki, and the study was approved by the Institutional Review Boards of the Ethics Committee of Xiangya Hospital, Central South University.

### PET Study

According to a previously reported method ^54^, positron emission tomography/computed tomography (PET/CT) was performed on the Family 2 II-4 using ^11^C-2β-carbomethoxy-3β-(4-fluorophenyl) tropane (^11^C-CFT) tracer. Before the PET/CT imaging, the patient discontinued the drug intake for 2 days to avoid the potential effect of anti-PD drugs. Brain PET imaging was performed at one hour after intravenous injection of ^11^C-CFT. The regions of interest in each hemisphere were identified and drawn on the caudate nucleus, putamen, and cerebellum.

### SNP genotyping and homozygosity mapping

DNA samples of Family 1 (AR-003), Family 2 (AR-005) and Family 4 underwent genome-wide SNP array genotyping. Genome-wide genotyping was performed with the Illumina Human Omni ZhongHua-8 Bead Chip arrays. Homozygosity mapping was performed with PLINK (http://pngu.mgh.harvard.edu/purcell/plink/) for the identification of regions of homozygosity in affected individuals, and the minimum length for homozygous runs was set to 2 Mb.

### Whole exome sequencing

Exome data were obtained from the 156 cases with ARPD, 1,498 cases of sporadic EOPD, and 1258 matched control subjects. As previous reported ^26^, whole-exome DNA was capture using the SureSelect Human All Exon Kit V5 or V6 (Agilent) and high-throughput sequencing was conducted using the Illumina X10 with a coverage more than 100 X. Burrow-Wheeler Aligner was implemented to align Paired-end sequence reads onto the reference human genome (UCSC hg19). The Picard tool (http://broadinstitute.github.io/picard/) was used to remove duplicate reads, generate the converse format, and index the sequencing data. Base quality-score recalibration, local realignments around possible insertions/deletions (indels), variant calling, and filtering were performed with the Genome Analysis Toolkit (GATK) ^55^. ANNOVAR ^56,57^ was used to annotate single nucleotide variants and insertions/deletions with RefSeq (UCSC hg19), such as gene regions, amino acid alterations, functional effects, and allele frequencies in East Asian population from GnomAD database and ExAC database. Mutations of previously reported PD causative genes were excluded. The minor allele frequency of the variants was limited to 0.01 for the above population database. Only predicted damaging missense and loss-of-function variants (nonsense variants, frameshift indels, and splicing-site variants) were included.

### Sanger sequencing

Potential mutations were confirmed by Sanger sequencing and were shown to segregate with the phenotype. Mutation analysis of *DAGLB* in another 500 matched control cohort was done by direct sequencing (GenBank, NM_139179.4 and NP_631918.3). Genomic DNAs from individuals were amplified by PCR with oligonucleotide primers complementary to flanking intronic sequences. Samples were run and analyzed on an ABI PRISM 3130 genetic analyzer (Applied Biosystems).

### Detection and validation of CNVs in *DAGLB*

The detection of copy-number variant (CNV) in *DAGLB* from WES data in our ARPD and sporadic EOPD cohorts was performed with the eXome-Hidden Markov Model (XHMM) software, which uses principal component analysis normalization and a hidden Markov model to detect and genotype CNVs from normalized read-depth data from targeted sequencing experiments. To further analyze the detailed structure variants of *DAGLB* in our patients identified by the WES CNV analysis, we used the Oxford Nanopore platform to sequence the same individuals (Family 3, AR-075). As previous reported^58,59^, large insert-size libraries were created according to the manufacturer recommended protocols (Oxford Nanopore). Libraries were sequenced on R9.4.1 flow cells using PromethION. NGMLR and Sniffles were used to analyze structural variations. All reads were aligned to the human reference genome (hg19) using NGMLR (ngmlr 0.2.7), and structural variation calls were detected by Sniffles. Candidate structural variations were subjected to manual examination and further validation. Sanger sequencing of the PCR product of the breakpoints was performed using standard protocols.

### Human skin fibroblast culture

Human dermal fibroblasts were derived from skin biopsies from affected individuals and age- and sex-matched non-neurological controls, through standard techniques ^60^. Fibroblasts were cultured in Dulbecco’s modified Eagle’s medium (ThermoFisher) supplemented with 10% fetal bovine serum, penicillin, and streptomycin (Gibco). Cells were grown in 5% CO2 at 37 °C in a humidified incubator. To further investigate the protein stability of DAGLB in human dermal fibroblasts, cells were treated with proteasome inhibitor MG132 (Sigma, M7449) at 10 μM for 24 hours before the cells were collected. DMSO was used as a control vehicle.

### Quantitative RT–PCR (qRT–PCR)

Total RNA was extracted with RNeasy kit (QIAGEN), and first-strand cDNA synthesis was performed with a SuperScript III First-Strand Synthesis system (Invitrogen). Real-time Taqman PCR was performed on an ABI 7900HT with TaqMan Gene Expression Assays (Applied Biosystems, Life Technologies, Carlsbad, CA) for human *DAGLB* exon 9-10 (Hs00373700_m1). Results were normalized to GAPDH. Experiments were performed with triplicate experimental samples and controls, and fold increases were calculated using the comparative threshold cycle method.

### Western blotting

Cells were lysed in radioimmunoprecipitation assay (RIPA) buffer containing protease and phosphatase inhibitor cocktails and sonicated for 1 min with the Bioruptor sonication device (Diagenode). Cell lysates were centrifuged at 13,000 × g for 10 min at 4 °C, and the supernatant was collected for protein quantification (Pierce BCA Protein Assay Kit). Each sample contained 20 μg of proteins and was mixed with Bolt LDS Sample Buffer and Sample Reducing Agent (ThermoFisher) and heated at 70 °C for 10 min. The prepared protein extracts were size fractioned by 4 to 12% NuPAGE Bis-Tris gel electrophoresis (Invitrogen) using MES running buffer (Invitrogen). After transfer to the nitrocellulose membranes using Transfer Cell (Bio-Rad), the membranes were blocked with Odyssey Blocking Buffer (LI-COR) and probed overnight with the appropriate dilutions of the primary antibodies. The antibodies used for western blot analysis included Rabbit monoclonal anti-DAGLB (Cell Signaling, 12574S, 1:500), Rabbit polyclonal anti-DAGLA (Frontier Institute co. ltd, DGLa-Rb-Af380, 1:250), Rabbit monoclonal anti-HA-Tag (Cell Signaling, 3724S, 1:500), Rabbit monoclonal anti-Cre Recombinase (Cell Signaling, 12830S, 1:500), Mouse monoclonal anti-GAPDH (Sigma-Aldrich, G8795, 1:5000) and Mouse monoclonal anti-β-actin (Sigma-Aldrich, A2228, 1:5000). Incubation with the IRDye-labeled secondary antibodies (LI-COR, 1:10000) was performed for 1 hour at room temperature. The protein bands of interest were visualized with Odyssey CLx Infrared Imaging Studio. The band intensity was quantified using ImageJ.

### Mouse work

All mouse studies were in accordance with the guidelines approved by Institutional Animal Care and Use Committees (IACUC) of the National Institute on Aging (NIA), NIH. The wild-type C57BL/6J (#000664), DAT^IRESCre^ (#006660), Ai95 (RCL-GCaMP6f) (#028865), and Ai9 (RCL-tdT) (#007909) mice were purchased from the Jackson laboratory. *Cnr1*^loxP/loxP^ mice ^38^ were generously provided by Dr. Josephine M. Egan of NIA. Mice were housed in a twelve-hour-light/twelve-hour-dark cycle and were fed water and regular diet ad libitum. All the behavioral tasks were performed during the light cycles. The genotype, gender and age of mice were indicated in the figure legends.

### RNA *in situ* hybridization with RNAscope

RNAscope (Advanced Cell Diagnostics, ACD) was performed according to the manufacturer’s instructions on fresh frozen tissue sections. The sample preparation and pretreatment were conducted according to the instructions of RNAscope Multiplex Fluorescent Reagent Kit v2 user manual. RNAscope probes for *Daglb* (Cat No. 497801-C1) and *Th* (Cat No. 317621-C2) were purchased from ACD and used according to the company’s online protocols. Fluorescent images were acquired using a laser scanning confocal microscope LSM 780 (Zeiss).

### RNA-sequencing

For the RNA sequencing of nigrostriatal DANs, we used LCM to isolate GFP-positive DANs in the SNc region of *Pitx3*^+/IRES2-tTA^ (JAX#021962)/pTRE-H2BGFP (JAX#005104) double transgenic mice as described previously ^61^. The animals were anesthetized with CO2 followed by decapitation at one year old. The brains were rapidly dissected and frozen in dry ice. The frozen brains were sectioned at 30µm thickness by a cryostat onto a PAN membrane frame slide (Applied Biosystems, Foster City, CA) and stored at -80°C until LCM performance. By an ArturusXT micro-dissection system with fluorescent illumination (Applied Biosystems), the GFP-positive cells in the SNc region were selected and then captured onto LCM Macro Caps (Applied Biosystems) at the following working parameters: spot size, 7-25µm; power, 50 – 70mW; duration, 20-40µs. The total RNA was extracted and purified with the PicoPure Isolation kit (Applied Biosystems) and genomic DNA was cleaned-up by RNase free DNase (Qiagen) after the protocols provided by the manufacturers. The RNA was quantified using a NanoDrop spectrophotometer (ThermoFisher) and the RNA integrity was measured using the Bioanalyzer RNA 6000 pico assay (Agilent). The cDNA libraries were generated from the purified RNA using TruSeq Stranded Total RNA LT library preparation kit (Illumina) according to the manufacturer’s instructions. The libraries were then qualified using the Bioanalyzer DNA 1000 assay (Agilent) and sequenced with Illumina HiSeq 2000. The standard Illumina pipeline was used to generate Fastq files. The Ensembl annotated transcript abundance was quantified using Salmon in a non-alignment-based mode, and gene level counts were estimated using Tximport package (Bioconductor). The counts for the resulting genes were then normalized using a variance-stabilizing transformation. For the RNA sequencing of striatal tissues, the dorsal striatal of 3-month-old C57BL/6J mice were dissected and subjected to RNA extraction, sequencing and data analysis as described recently ^62^.

### AAV-*Daglb* KD gene targeting vector construction, validation, and packaging

#### Vector construction

All plasmids were constructed using standard recombinant DNA cloning techniques. The PX601-AAV-CMV::NLS-SaCas9-NLS-3xHA-bGHpA;U6::BsaI-sgRNA plasmid was a gift from Feng Zhang (Addgene plasmid # 61591) ^28^. The *Daglb* sgRNA oligos were designed with Benchling (https://benchling.com) and subcloned into the PX601-AAV-CMV::NLS-SaCas9-NLS-3xHA-bGHpA;U6::BsaI-sgRNA vector. To construct a single AAV vector harboring both Cre-dependent SaCas9 transgene and constitutively expressed sgRNA expression cassette, the Magneto2.0-sNRPpA element of pAAV-CMV-DIO-Magneto2.0-sNRPpA expression vector, a gift from Ali Guler (Addgene plasmid # 74307)^63^, was replaced by the SaCas9-NLS-3xHA-bGHpA;U6::BsaI-sgRNA DNA fragment.

#### Vector validation

Neuro-2a (N2a) cell lines were maintained in Dulbecco’s modified Eagle’s medium (DMEM) supplemented with 10% FBS (HyClone), 2mM GlutaMAX (Life Technologies), 100U/ml penicillin, and 100 mg streptomycin at 37°C with 5% CO2 incubation. Cells were co-transfected with pAAV-Cre-GFP and pAAV-CMV-DIO-SaCas9-NLS-3xHA-bGHpA;U6::BsaI-sgRNA plasmids or pAAV-EF1a-DIO-mCherry plasmids (Addgene plasmid #50462) at the ratio of 1:3 using X-tremeGENE HP DNA Transfection Reagent (Roche) following the manufacturer’s recommended protocol. Cells were harvested for PCR-based identification of mutations caused by genome editing using the Guide-it Mutation Detection Kit (Cat. No. 631443), and immunoblotting were performed to analyze the DAGLB protein levels.

#### AAV packaging

The packaging was carried out by a commercial source (Vigene Biosciences Inc.) and the resulting AAVs had titers of 1.0 × 10^13^ to 2.0 × 10^14^ genome copies per ml.

#### Stereotactic injection

The stereotactic survival surgery was performed as previously described^7^. 500 nL of AAVs with titers 8.0 × 10^13^ genome copies per ml were loaded into 2 μL Neuros Syringes (Hamilton) and were injected into brain areas at chosen coordinates. The coordinates based on Bregma coordinate for SNc are AP -3.1 mm, ML: ± 1.5 mm, DV -3.9 mm.

### Primary neuronal culture and viral infection

Mouse primary neuronal cultures were prepared from the cortices of embryonic day 16.5 embryos. Briefly, cortices were dissected in cold Hank’s balanced salt solution, and incubated with 0.025% trypsin for 20 min at 37°C. The digested tissue was triturated into single cells using glass Pasteur pipettes and filtering through 70 µm nylon cell strainer. The cells were seeded in plated in Biocoat Poly-D-Lysine Cellware plate and maintained in neurobasal medium supplemented with 2% B27 and 2 mM GlutaMax at 37°C in 5% CO2 humidified incubator. Cells at 4 days in vitro (DIV) were infected with AAV DJ-Cre-GFP and AAV DJ-CMV-DIO-SaCas9-NLS-3xHA-bGHpA;U6::BsaI-sgRNA at the ratio of 1:3, and the cells were harvested 7 days after infection for western blot analyses.

### Behavioral tests

#### Rotarod motor skill learning test

Mice were placed onto a rotating rod with auto acceleration from 4 to 40 rpm in 5 min (Panlab). The duration that each mouse was able to stay on the rotating rod in each trial was recorded as the latency to fall. The standard motor learning task was performed as ten trials per day for six consecutive days as described previously ^7^.

#### Open-field test

The ambulatory, rearing and fine movements of mice were measured with the Flex-Field activity system (San Diego Instruments). Flex-Field software was used to trace and quantify mouse movement in the unit as the number of beam breaks per 30 min as previously described ^64^.

#### Gait analysis

The Free Walk Scan system (CleverSys Inc) was used for gait analysis as described before ^7^. Briefly, mice were allowed to move freely in a 40 cm × 40 cm × 30 cm (length × width × height) chamber. A high-speed camera below a clear bottom plate was used to capture mouse movement for 5 min in the red light. Videos were analyzed using FreewalkScanTM2.0 software (CleverSys Inc) for various characteristic parameters of gait including stride length and stance/swing time of each paw.

### Histology, immunohistochemistry, and light microscopy

Mice were anesthetized with ketamine and then transcardially perfused with 4% PFA/PBS solution. Brains were isolated, post-fixed in 4% PFA overnight, and then submerged in 30% sucrose for 72 hour at 4 °C for later sectioning. Series of 40 μm sections were collected using a cryostat (Leica Biosystems). Sections were blocked in 10% normal donkey serum, 1% bovine serum albumin, 0.3% Triton X-100, and PBS solution for overnight at 4 °C. The sections were then incubated with the primary antibodies over one to two nights at 4 °C. The antibodies used for immunostaining included rat monoclonal anti-DRD1 (Sigma-Aldrich, D2944, 1:500), mouse monoclonal anti-CB1 (Synaptic systems, 258011, 1:500), rabbit monoclonal anti-TH (Pel-Freez Biologicals, P40101, 1:2500), mouse monoclonal anti-TH (ImmunoStar, 22941, 1:1000), chicken polyclonal anti-TH (Aves Labs, TYH, 1:500), chicken polyclonal anti-GFP (Aves Labs, GFP-1020, 1:1000), rabbit polyclonal anti-RFP (Rockland, 600-401-379, 1:1000), rabbit monoclonal anti-HA-Tag (Cell Signaling, 3724S, 1:100) and guinea pig polyclonal anti-NeuN (Synaptic systems, 266 004, 1:1000). Sections were then washed three times in PBS before being incubated in the secondary antibody solutions with Alexa Fluor 488, 546, or 633-conjugated secondary antibodies (1:500, Invitrogen) at 4 °C for overnight. Following three washes in PBS, sections were mounted onto subbed slides, and coverslipped with mounting media (ProLong^®^ Gold Antifade Mountant, Life technology). The stained sections were imaged using a laser scanning confocal microscope (LSM 780, Zeiss). The paired images in the figures were collected at the same gain and offset settings.

### Stereology

According to the mouse brain in stereotaxic coordinates, a series of coronal sections across the midbrain (40 μm per section, every fourth section from Bregma − 2.54 to − 4.24 mm, ten sections per case) were processed for TH immunohistochemistry and finally visualized using a laser scanning confocal microscope (LSM 780, Zeiss). The images were captured as a single optic layer under 20 × objective lens. TH-positive neurons in SNc were assessed using the fractionator function of Stereo Investigator 10 (MBF Bioscience) as described previously ^61^. Five mice were used per group. Counters were blinded to the genotypes of the samples.

### 2-AG measurement with liquid chromatography-tandem mass spectrometry

Endocannabinoids were extracted from the SNc of 3 to 4-month-old mice and quantified by LC-MS/MS as previously described ^65^. In brief, the fresh brain tissues were sliced at 500 μm thickness and frozen immediately in liquid nitrogen. The samples were taken by punch technique then kept on dry ice or at -80°C. Tissue samples from individual mice were homogenized in 80-300 μl of Tris buffer (pH 8.0) and the protein concentrations were determined by Bradford assay. Ice-cold of methanol/Tris buffer (50 mM, pH 8.0) solution was added to each homogenate (1:1, vol/vol). 200 ng [^2^H_5_] of arachidonoyl glycerol ([^2^H_5_]2-AG) were used as internal standard. The homogenates were extracted three times with CHCl_3_: methanol (2:1, vol/vol), dried under nitrogen and reconstituted with methanol after precipitating proteins with ice-cold acetone. The dried samples were reconstituted in 50 μl of ice-cold methanol, and 2 μl of which were analyzed with liquid chromatography in line mass spectrometry. The LC-MS/MS analyses were conducted on an Agilent 6410 triple quadrupole mass spectrometer (Agilent Technologies) coupled to an Agilent 1200 LC system. Analytes were separated using a Zorbax SB-C18 rapid-resolution HT column. Gradient elution mobile phases consisted of 0.1% formic acid in water (phase A) and 0.1% formic acid in methanol (phase B). Gradient elution (250 mL/min) was initiated and held at 10% B for 0.5 min, followed by a linear increase to 85% B at 1 min and maintained until 12.5 min, then increased linearly to 100% B at 13 min and maintained until 14.5 min. The mass spectrometer was set for electrospray ionization operated in positive ion mode. The source parameters were as follows: capillary voltage, 4,000 V; gas temperature, 350°C drying gas, 10 L/min; nitrogen was used as the nebulizing gas. Collision-induced dissociation was performed using nitrogen. Level of each compound was analyzed by multiple reactions monitoring. The molecular ion and fragment for each compound were measured as follows: m/z 348.3/91.1 for [^2^H_5_]2-AG and m/z 379.3/91.1 for 2-AG. Analytes were quantified by using Mass-Hunter Workstation LC/QQQ Acquisition and MassHunter Workstation Quantitative Analysis software (Agilent Technologies). Levels of AEA and 2-AG in the samples were measured against standard curves.

### *In vivo* fiber photometry

A custom-built dual color fiber photometry system ^31^ was used for *in vivo* measurement of eCB2.0, DA2m, GCaMP6f, and tdTomato fluorescent signals. For imaging the eCB2.0 or DA2m signals, 600 nl AAV9-hsyn-eCB2.0 (2.3×10^12^ GC/ml, Vigene Biosciences) or AAV9-hsyn-DA2m (4.67×10^11^ GC/ml, Vigene Biosciences) AAVs were mixed with 200 nl AAV9-hsyn-tdTomato (1.38×10^12^ GC/ml, Vigene Biosciences) AAVs and stereotactically injected in the dorsal striatum (coordinates: AP+0.5mm, ML+2.4mm, DV -2.5mm; AP+1.5mm, ML+1.8mm, DV-3.0mm) of 3 to 4-month-old wild-type C57BL/6J (JAX#000644) or DAT^IRES*cre*^ (JAX#006660) mice. Four weeks after viral injection, an optical probe (200 μm core and 0.22 NA) was implanted with the tips sitting in the SNr areas (coordinates: AP-3.16 mm, ML+1.4 mm, DV -4.4 mm) for imaging the eCB2.0 or DA2m signals. The animals were allowed to recover for at least one week after the fiber implantation surgery before the fiber photometry measurement. The fluorescence signals were acquired using 49 ms integration time and were triggered by 20 Hz transistor-transistor logic pulses from an output pulse generator. The eCB2.0, DA2m or GCaMP6f fluorescence signals were calculated by total photo counts between 500 nm and 540 nm. The tdTomato fluorescence signals were calculated by total photon counts between 575 nm and 650 nm. The measured emission spectra of eCB2.0, DA2m, GCaMP6f and tdTomato signals were fitted using a linear unmixing algorithm (https://www.niehs.nih.gov/research/atniehs/labs/ln/pi/iv/tools/index.cfm). The coefficients of eCB2.0, DA2m, GCaMP6f and tdTomato signals generated by the unmixing algorithm were used to represent the fluorescence intensities of eCB2.0, DA2m, GCaMP6f and tdTomato, respectively. To correct for movement-induced artifacts, the ratios of eCB2.0, DA2m or GCaMP6f signal intensities against the corresponding tdTomato signal intensities were used to represent the final normalized signal intensities.

For the JZL184 experiments, fiber photometry recordings were conducted in free-moving animals for 20 min before drug administration to measure the baseline fluorescent intensities and then for 60 min or 120 min after drug treatment. The average baseline signals were calculated as F_B_. The instant signals at different time point after drug treatment were calculated as F_I_. The alterations of signal intensities at different time points were calculated as ΔF/F=(F_I_-F_B_)/F_B._

The rotarod motor skill learning and fiber photometry recoding experiments were performed as reported previously^34^. Briefly, in each trial the mice were put on a rotatable rod (EZRod, Omnitech Electronics) starting at 4 rpm constant speed for 30 sec and then steadily accelerated from 4 to 40 rpm in 5min, while the fiber photometry recording was performed at the same time. 10 trials and recordings were carried out each day for six continuous days. F_B,_ the baseline signal intensity of the first trial of each day, is the average signal intensities at 4 rpm for the first 30 sec. F_I_ represents the average signal intensity during each trial. The alterations of signal intensities at different trials were calculated as ΔF/F=(F_I_-F_B_)/F_B._

### Statistical analyses

All the data were analyzed by Prism 8 software (Graphpad). Data were presented as mean ± SEM or mean ± SD. N represents animal numbers and is indicated in the figure legends. Statistical significance was determined by comparing means of different groups using *t* test or ANOVA followed by post hoc tests.

## Acknowledgements

This work is supported in part by the National Key Plan for Scientific Research and Development of China grants (BT, Grant No. 2016YFC1306000), the National Natural Science Foundation of China (BT, Grant No. 81430023), and the Intramural Research Programs of National Institute on Aging, NIH (HC, ZIA AG000944, AG000928), National Institute of Alcoholism and Alcohol Abuse (DML, ZIA AA000416; AS, K99/R00 AA025991), and National Institute of Environmental Health Sciences (GC, ZIA ES103310). We thank NIMH rodent behavioral core for assisting in behavioral tests, Dr. Josephine M. Egan of NIA for providing the *Cnr1*^loxP/loxP^ mice and members of Cai lab for their suggestions and technical assistance. Drs. Zhenhua Liu and Nannan Yang used to be participants in the NIH Graduate Partnership Program and graduate students at Central South University. Dr. Wotu Tian was a participant in the NIH Graduate Partnership Program and a graduate student at Shanghai Jiao Tong University School of Medicine. Dr. Jie Dong used to be a participant in the NIH Graduate Partnership Program and a graduate student at Dalian Medical University. We are indebted to the participation of the patients and their family members in this study.

## Author contributions

B.T. designed and supervised the human genetics study. H.C. conceived, designed and supervised the mouse experiments. H.C. wrote the manuscript and prepared the figures with inputs from all authors. Z.L. performed human genetics study, biochemistry, histology and mouse behavioral experiments, and prepared figures and tables of genetics data. N.Y. performed fiber photometry experiments and data analyses. J.G., J.M., P.C., H.S. and T.W. contributed to human genetic study. J.T. and Z. Z. contributed to biochemistry study. W.T., S.C., S.H., J.K., and J.W. contributed to behavior tests, histology, and data analyses. A.S., D.L., J.D., W.L., J.Z. and G.C. contributed to fiber photometry, histology, and data analyses. L.C. performed stereotactic surgery. L.S., C.X. and J.H.D. contributed to RNA-sequencing and data analyses. Y.L., A.D., and K.H. provided eCB2.0 and DA2m sensors. R.C. performed endocannabinoids measurements. All authors read and approved the final manuscript.

