## Supplementary material for "Deficiency in endocannabinoid synthase *DAGLB* contributes to Parkinson’s disease and dopaminergic neuron dysfunction": Table S1-S5, Fig. S1-S14

**Supplementary Clinical description, Tables and Figures.**

#### Clinical description

##### Family 1

The family consists of healthy parents who were first-degree relatives, two affected, and three unaffected siblings. The birth, early milestones and development of these two affected females were normal, but they developed the motor symptoms, tremor and bradykinesia during the fourth decade of life. The first patient (II-3) developed a mild right leg tremor at the age of 36. One year later, she had resting tremor in the left lower limb, and developed a slightly masked face, a stiff-legged gait and slowness of movements. Brain magnetic resonance imaging (MRI) was reported to be normal at age 38 and her symptoms improved after introduction of Levodopa-Benserazide. She developed hallucination with the introduction of Piribedil which disappeared one month later. Neurological examination performed at the age of 41 showed hyposmia, depression, urinary urgency, sleep disturbance, rapid eye movement sleep behavior disorder (RBD), and cognitive function was grossly normal. The deep tendon reflexes were normal and pyramidal signs were absent. A new MRI was negative and other investigations were normal. The second patient (II-4) in this family developed a mild right arm tremor at the age of 34 years. Subsequently, she had stiffness and slowed movements in the right hand. She was treated with Levodopa-Benserazide and Piribedil at this time with improvement in both tremor and bradykinesia. She deteriorated gradually and presented bilateral resting tremor, rigidity and bradykinesia in the next two years. She presented hyposmia, depression, urinary urgency, constipation, sleep disturbance and RBD without cognitive dysfunction on our neurological examination at the age of 39. Tendon reflexes were normal and pyramidal signs were absent. Cranial MRI was unremarkable and other laboratory tests were normal. Both patients in this family showed a very similar phenotype and their clinical course was typical for PD, including good initial response to levodopa with marked contrast in motor function during on versus off state, and on-state dyskinesia.

**Family 2**

The female patient was born in a consanguineous family with healthy parents and three unaffected siblings. She had an uneventful birth and normal development until adulthood. At the age of 37, she developed mild slowness of movement in the right arm. Subsequently, she experienced difficulty walking with dragging of the right foot. Over the years, the motor disorder progressed slowly. She presented with bradykinesia, rigidity and mild resting tremor in subsequent years. Hypomimia, depression, urinary urgency, RBD and sleep disturbance were presented. After 13 years of disease, there were no signs of dementia. Deep tendon reflexes and eye movements were normal. Brain MRI was unremarkable, whereas $^{11}$C-2β-carbomethoxy-3β-(4-fluorophenyl) tropane ($^{11}$C-CFT) positron emission tomography (PET) imaging revealed a graded and asymmetrical reduction in dopamine transporter binding in the putamen which was compatible with PD (**Supplementary Fig. S6**). This patient responded well to dopaminergic therapies but developed severe dyskinesia after levodopa-benserazide treatment. She received deep brain stimulation after ten years of disease, which improved her motor symptoms and dyskinesia.

**Family 3**

This family included two female patients suffering recessive early-onset forms of PD. The proband patient II-2 started suffering from right-hand tremor at the age of 37. One year later, she developed tremor in the left limb. Over the subsequent year, she developed slowly progressive bradykinesia, fatigue, leg stiffness, and gait impairment. Levodopa/benserazide therapy, initiated within two years of the onset of symptoms, yielded clear benefits to her motor disturbances. She also received left side posteroventral pallidotomy after five years of disease, which improved her symptoms. On physical examination 13 years after disease onset, she had severe parkinsonism and was wheelchair-bound. She had severe axial and forelimb rigidity, more marked on the right side. The second case patient II-1 from this family developed resting tremor in the left leg at age 39. Then she presented resting tremor in the left hand and slowed movements and stiffness in the left limb. Brain MRI was reported to be normal and the patient was treated with levodopa with very good response at the age of 40. Two years later, she developed slowness of movements and tremor in the right limb. She presented severe depression and cognitive decline (The score of the Mini-Mental State Examination (MMSE) was 15) on the neurological examination 14 years after the onset of the disease. The deep tendon reflexes were symmetric and normal, and the Babinski sign was absent. Motor complications with treatment including treatment wearing off and On-off phenomenon were present, but no dyskinesia was observed in these two patients on our neurological examination. The disease symptoms of both patients are typical for PD of long disease duration.

**Family 4**

A male patient with sporadic PD developed slowly progressive symptoms starting at the age of 35. He developed all cardinal signs of PD including resting tremor, rigidity and bradykinesia in the subsequent years. No atypical clinical features were presented. Neurologic examination showed he had hyposmia and mild depression. Brain MRI was negative and other blood chemistry was normal. This patient responded well to dopaminergic therapies.

#### Supplementary Table 1

**Table S1. Homozygous segments larger than 2Mb identified in both affected sibling in Family 1 by homozygosity mapping through genome-wide SNP genotyping**

| No. | Chromosome | Start | End | Length (Kb) | SNP number | SNP density (SNP per kb) | Proportion of sites homozygous | Proportion of sites heterozygous |
|---|---|---|---|---|---|---|---|---|
| 1 | chr3 | 9244585 | 15423557 | 6179 | 2669 | 2.315 | 0.993 | 0 |
| 2 | chr5 | 24837962 | 30738438 | 5900 | 1396 | 4.227 | 0.993 | 0 |
| 3 | chr6 | 100815 | 2631241 | 2530 | 1151 | 2.198 | 0.995 | 0.003 |
| 4 | chr7 | 1733102 | 8721760 | 6989 | 2674 | 2.614 | 0.999 | 0 |
| 5 | chr17 | 30861865 | 34353274 | 3491 | 1419 | 2.46 | 0.994 | 0.001 |

#### Supplementary Table 2

##### Table S1. Whole-exome sequencing in three PD families

| Family | Family 1 | | Family 2 | Family 4 |
|---|---|---|---|---|
| Family member | II-3 | II-4 | II-4 | II-1 |
| Total clean reads | 87,161,148 | 82,704,614 | 80,568,752 | 63228822 |
| Total clean data (Mb) | 10849.79 | 10269.50 | 10021.30 | 9368.32 |
| Mapped rate (%) | 99.89 | 99.84 | 99.91 | 99.74 |
| Mean depth | 137.90 | 130.75 | 128.22 | 111.60 |
| Coverage≥10X | 99.0% | 97.8% | 99.1% | 98.3% |
| Coverage≥20X | 97.5% | 95.1% | 97.4% | 99.5% |
| Total variants | 38837 | 37991 | 38454 | 39864 |
| Nonsynonymous variants in exon or splicing (±2bp) region | 11814 | 11357 | 11770 | 12273 |
| MAF＜0.01 in East Asian population in GnomAD exome, GnomAD genome, and ExAC | 547 | 557 | 577 | 597 |
| Deleterious predicted by Reve | 114 | 114 | 119 | 126 |
| Homozygous variants in affected individuals | *DAGLB* | | *DAGLB, OPLAH, HABP2, SSBP4* | *DAGLB* |
| Compound heterozygous variants in affected individuals | *FLG, ZNF806, PRIM2, TIMELESS, KNTC1* | | *LRP8, FLG, YY1AP2, ZNF806, FRG1, PRIM2, PRSS1, TDG, ACOT4* | *FLG, ZNF806, TTLL3, PRIM2, TAS* |

#### Supplementary Table 3

**Table S3. Identified mutations in *DAGLB* and predictions of their pathogenicity**

| Family | Family 1 | Family 2 | Family 4 |
|---|---|---|---|
| Zygosity | homozygous | homozygous | homozygous |
| Chromosome Postion[a] | ch7: 6449668 | ch7: 6464435 | ch7: 6474600 |
| cDNA alteration[b] | c.1821-2A>G | c.1088A>G | c.470dupC |
| Amino Acid Alteration | modify donor splice sites | p.D363G | p.L158Sfs*17 |
| MutationTaster[c] | Deleterious | Deleterious | Deleterious |
| CADD[c] | Deleterious | Deleterious | Deleterious |
| Reve[c] | Deleterious | Deleterious | Deleterious |
| gnomAD_exome_EAS[d] | absent | absent | absent |
| gnomAD_genome_EAS[d] | absent | absent | absent |
| ExAC_EAS[d] | absent | absent | absent |
| Chinese control cohort 1 (n=1,258, investigated by Whole-exome sequencing)[e] | absent | absent | absent |
| Chinese control cohort 2 (n=500, investigated by Sanger direct sequencing)[f] | absent | absent | absent |

[a]Position on Genome Reference Consortium human genome build 37 (GRCh37)

[b]Accession number for *DAGLB* is NM_139179

[c]Mutation prediction by MutationTaseter, CADD and Reve

[d]Frequency of the mutation in East Asian population (from genomAD exome, genomeAD genome, and ExAC databases) was calculated by mutated allele number/total allele number in parentheses

[e]Allele frequencies of these pathogenic mutations in Han Chinese controls consisted of 1,258 healthy individuals detected by Whole-exome sequencing

[f]Allele frequencies of these pathogenic mutations in Han Chinese controls consisted of 500 healthy individuals detected by Sanger direct sequencing

#### Supplementary Table 4

##### Table S4. Clinical characteristics of patients with biallelic *DAGLB* mutations

| | Family 1 II-3 | Family 1 II-4 | Family 2 II-4 | Family 4 II-1 | Family 3 II-1 | Family 3 II-2 |
|---|---|---|---|---|---|---|
| Mutation | c.1821-2A>G | c.1821-2A>G | c.1088A>G: p.D363G | c.469dupC: p.L158Sfs*17 | g.chr7:6,486,383-6,489,136 del | g.chr7:6,486,383-6,489,136 del |
| Gender | Female | Female | Female | Male | Female | Female |
| Age at onset, years | 36 | 34 | 38 | 35 | 39 | 37 |
| Age at examination, years | 40 | 38 | 49 | 41 | 43 | 50 |
| Disease duration, years | 5 | 5 | 11 | 6 | 7 | 13 |
| Symptoms at onset | Resting tremor | Resting tremor | Bradykinesia | Bradykinesia | Resting tremor | Resting tremor |
| Asymmetry at onset | + | + | + | + | + | + |
| Hoehn-Yahr stage (Off/On) | IV/II | IV/II | IV/II | III/II | III/II | V/III |
| Motor symptom | | | | | | |
| Bradykinesia | + | + | + | + | + | + |
| Resting tremor | + | + | + | + | + | + |
| Rigidity | + | + | + | + | + | + |
| Postural instability | + | + | + | + | + | + |
| UPDRS III (Off/On) | 56/37 | 74/40 | 76/33 | 50/28 | 52/30 | 76/58 |
| Nonmotor symptom | | | | | | |
| Hypomimia | + | + | + | + | + | + |
| Depression | + | + | + | + | + | + |
| Urinary urgency | + | + | + | + | - | + |
| Constipation | - | + | + | + | - | + |
| Cognitive decline | - | - | - | - | + | - |
| Hallucination | + | - | - | - | - | - |

| | | | | | | |
|---|---|---|---|---|---|---|
| Sleep disturbance | + | + | + | + | + | + |
| Freezing gait | + | + | + | + | + | + |
| RBD | - | + | + | - | + | - |
| Response to levodopa | + | + | + | + | + | + |
| Complications with treatment | | | | | | |
| Wearing off | + | + | + | + | + | + |
| On-off phenomenon | + | + | + | + | + | + |
| Dyskinesia | + | + | + | + | - | - |
| Surgical therapies | NA | NA | +* | NA | NA | +# |
| Brain MRI | - | - | - | - | - | - |
| Brain $^{11}C$-CFT PET | NA | NA | Abnormal* | NA | NA | NA |

+ = Present; - = Absent; NA=Not performed

* Deep brain stimulation

### Posteroventral pallidotomy

*Severe striatal uptake deficit, particularly at putamen level, as seen in Parkinson's disease

MRI=magnetic resonance imaging; PET=positron emission tomography; CFT=C-2β-carbomethoxy-3β-(4-fluorophenyl) tropane

#### Supplementary Table 5

**Table S5 List of primers used in this study**

Splicing analysis

| RT-PCR | Primer name | Sequence |
|---|---|---|
| c.1821-2A>G | RT-PCR Primer Forward | GATGTGATTCCCAGGCTCAG |
| | RT-PCR Primer Reverse | TCAGGCCACGTCCACACT |

Breakpoint PCR

| PCR | Primer name | Sequence |
|---|---|---|
| g.ch7:6,486,383-6,489,136del | Primer F1 | CCAAAACAAGGCAAGGTTCACT |
| | Primer R1 | CAAGTAGCCAGGACTACAGGTGC |
| | Primer F2 | TCTCGGTTCAACACGCAAGCCCCT |
| | Primer R2 | CTGCACCCTGCCTGGGACT |
| | Primer F3 | CCAAAACAAGGCAAGGTTCACT |
| | Primer R3 | GGGTCTCACTCTGCTACCCAGG |

*DAGLB* direct sequencing

| Primer name | Sequence |
|---|---|
| EXON1-F | GCAGACCTGCAATCGACTC |
| EXON1-R | CCACTTCTGTCACCGTCTCA |

| | |
|---|---|
| EXON2-F | GCCCTGTCCCCTTTTATTTC |
| EXON2-R | CGCCTGGTACAGGATTTCTT |
| EXON3-F | AAAGTCAGGAGGCGGGTG |
| EXON3-R | CGCAGAAACTAACTCCAATTTC |
| EXON4-F | CCTCGGTAACAGAACCCTCC |
| EXON4-R | CCACACACCCAAAGACACC |
| EXON5-F | CTGCCAGGAGCAGTCTTTTT |
| EXON5-R | GAGGGGAAAGGGGAATGA |
| EXON6-F | GAGAGTTCTATTCAACGAAGGAGC |
| EXON6-R | TACAGCGCATGTGACCAG |
| EXON7-F | AAAAGCCATTCCATGTCAGC |
| EXON7-R | TGGCCCCTCTGAAATAGTACAC |
| EXON8-F | CCCACTGTGTAGTGAGCGTG |
| EXON8-R | TCTGTCCTCACCATTTTCCC |
| EXON9-F | CGCTGGGTTTCCCTCTTTAG |
| EXON9-R | GCTCTGTGCAGTCCCTGAC |
| EXON10+11-F | GACGTGGCTGCTGATTCTG |
| EXON10+11-R | TACAGACACCCGCCAGCAT |
| EXON12+13-F | AACTGACGTTTCCCCTACCC |
| EXON12+13-R | TGATCAGATGGTGGAAGGAG |
| EXON14+15-F | GCATCTTTGCTGGAGTCTTCC |
| EXON14+15-R | GACCATGGAATTCTGTTCCC |

### Supplementary Figure 1

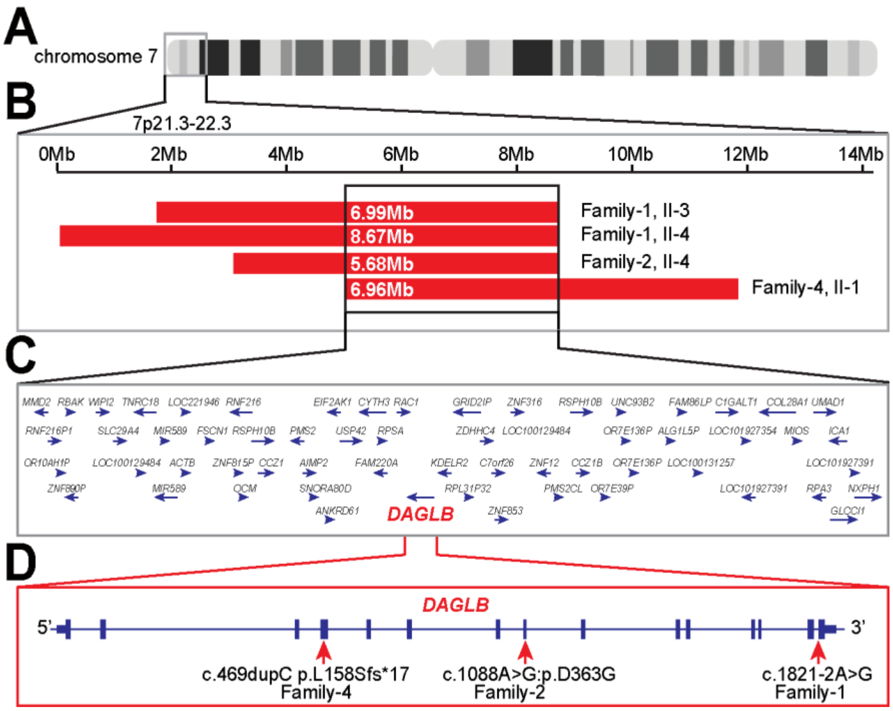


**Fig. S1 Molecular genetic findings in patients with *DAGLB* variants**. **(A)** Schematic of chromosome 7. **(B)** Homozygosity mapping of affected families showing the overlapping regions of homozygosity on 7q21.3-7q22.3 (GRCh37/hg19). Run of homozygosity regions for each case are shown as red boxes. Black lines indicate the minimal overlap region. **(C)** Genes located in the overlap homozygosity region. **(D)** Schematic of the exon-intron structure of *DAGLB* indicating the positions of the frameshift insertion, missense mutation, and splice-site variant.

#### Supplementary Figure 2

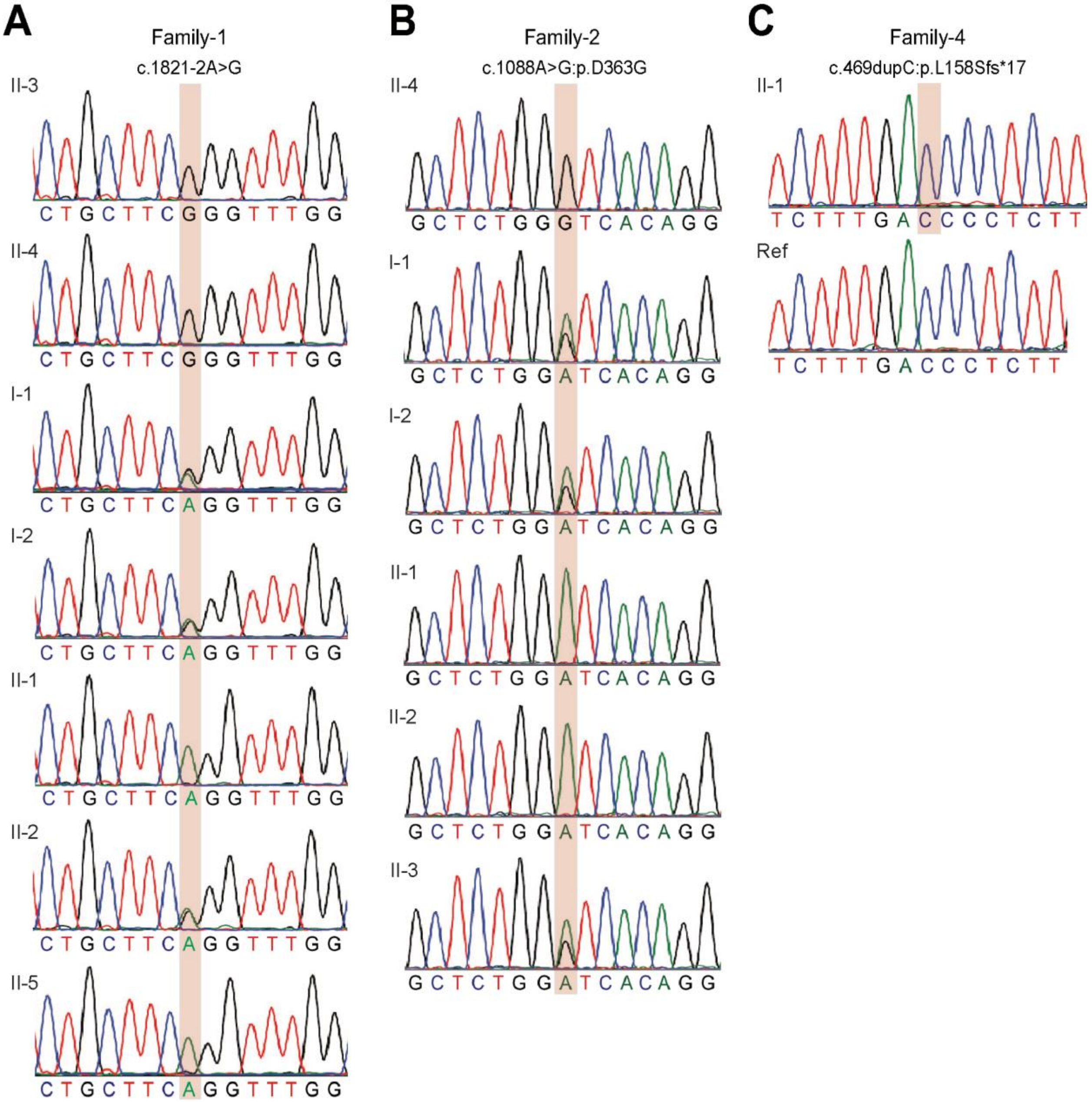


**Fig. S2 Sanger confirmation and segregation of the identified *DAGLB* variants.** Sequence chromatograms of patients, their parents and siblings from Family-1 **(A)**, Family-2 **(B)**, and Family-4 **(C)** (where available) are shown. Mutations were marked by pink box.

#### Supplementary Figure 3

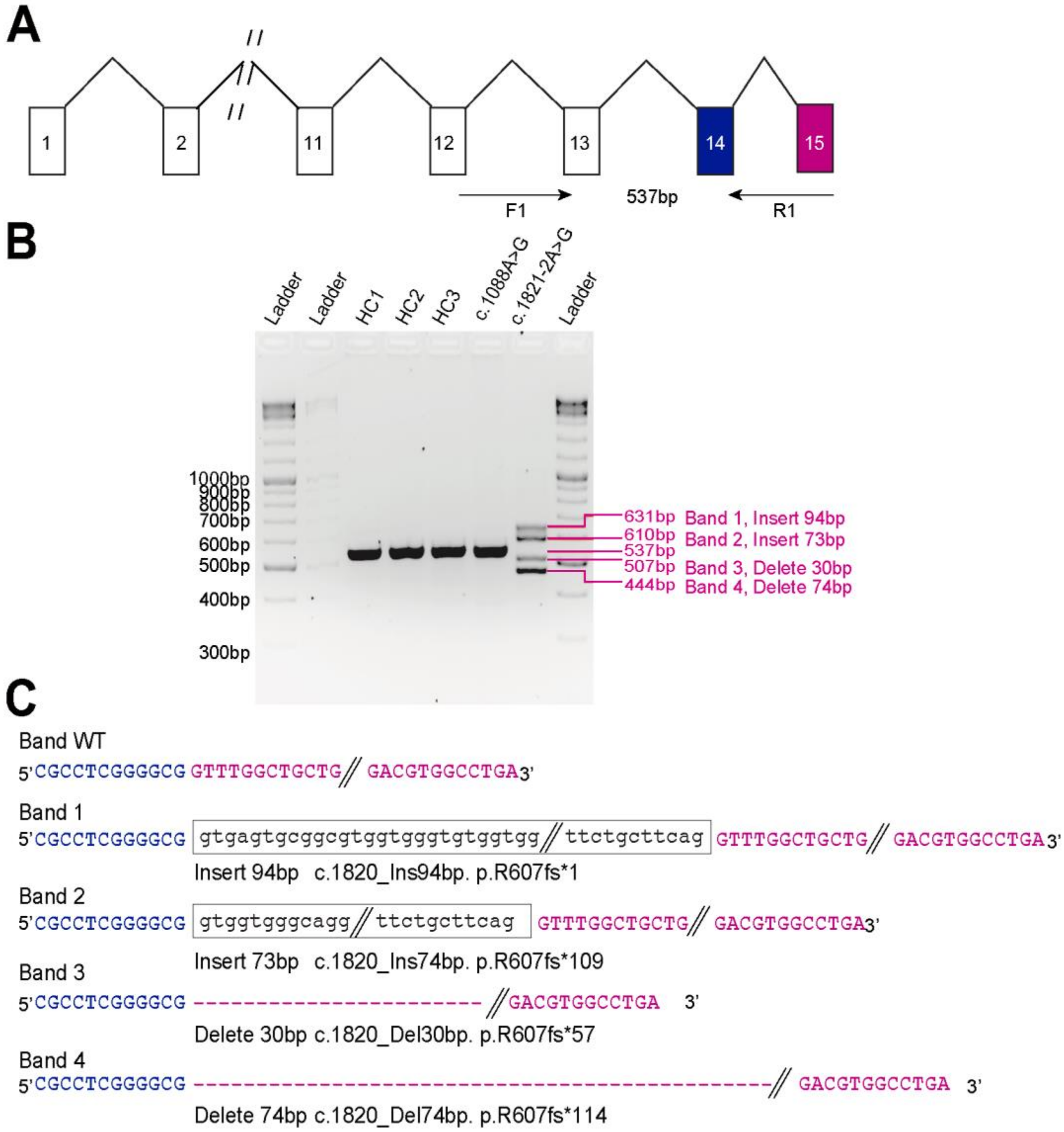


**Fig. S3 The c.1821-2A>G mutation resulted in aberrant splicing of *DAGLB*. (A)** A partial gene structure for *DAGLB*. Exons are boxed. The primer pair (F1 and R1) used for amplification are shown as arrows below the exon. **(B)** Fibroblast RNA samples from the affected individuals (Family 1 II-3, Family 2 II-4) and three health control subjects demonstrated four bands (631bp, 610bp, 507bp and 444bp) for c.1821-2A>G variant (Family 1 II-3) but only one for other subjects. (C) Gel purification, PCR and Sanger sequencing of the four bands from Family 1 II-3 demonstrate wild-type (WT) sequence and abnormal splicing sequence.

#### Supplementary Figure 4

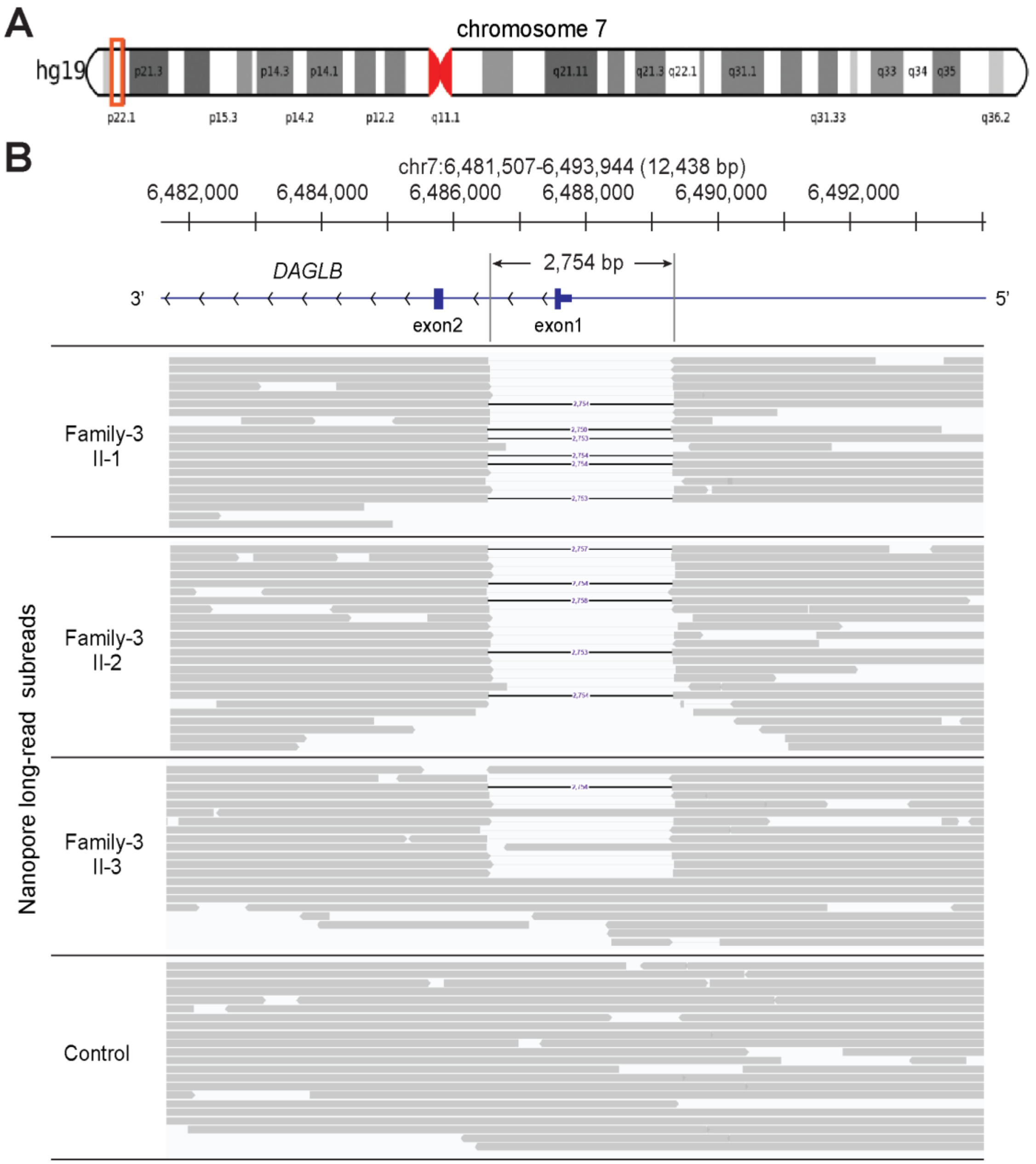


**Fig. S4** ***DAGLB*** **deletion detected by Nanopore long-read sequencing. (A)** Schematic of chromosome 7. **(B)** Nanopore long-read sequencing identifies a 2,754 bp deletion that includes the first coding exon of *DAGLB*. Nanopore long-read data were aligned to the human genome reference sequence (GRCh37/hg19). The site of deletion is shown by black connecting lines and subreads are shown by gray boxes. Nanopore long read subreads are shown for control, health sibling and the affected sisters from the Family 3. A homozygous 2,754 bp deletion involving *DAGLB* was supported by 20 or 24 Nanopore long reads without any read supporting the reference allele in the affected individuals (Family 3 II-1 and II-2) respectively, and 11 of 21 reads at the locus support the heterozygous deletion in the carrier sibling (Family 3 II-3).

#### Supplementary Figure 5

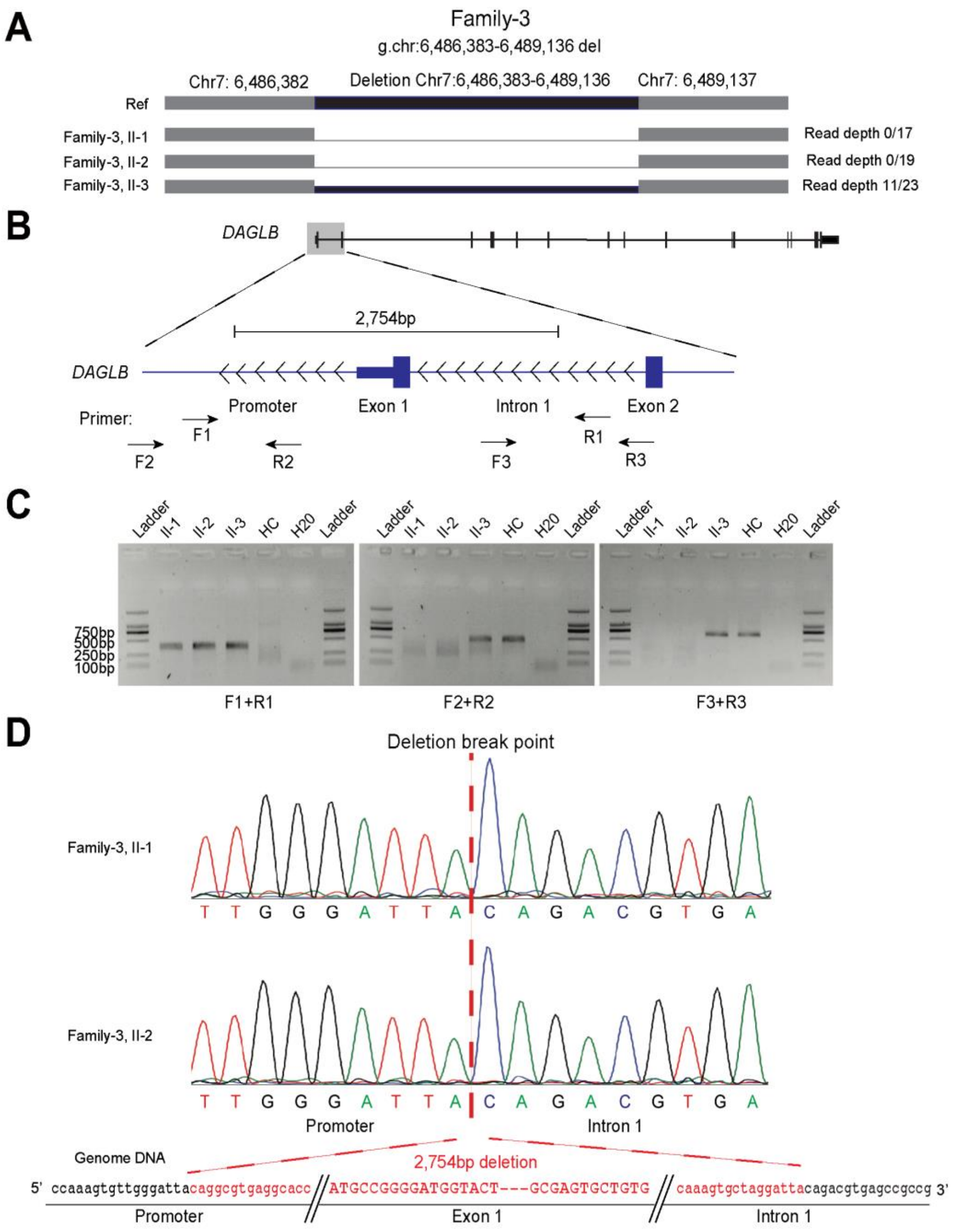


**Fig. S5 Characterization of a 2.7-kb deletion at the *DAGLB* locus. (A)** The high-quality Nanopore long-reads identify a 2,754 bp deletion (chr7:6,486,383-6,489,136) involving *DAGLB*. **(B)** Using positional information from Nanopore calls, PCR primers were designed to amplify the breakpoint junction. **(C)** The junction fragment was amplified when using DNA from the affected individuals (homozygous deletion) and the carrier sibling (heterozygous deletion), but not from control DNA. **(D)** This deletion variant was validated by Sanger sequencing, confirming the precise breakpoints identified by long reads sequencing.

#### Supplementary Figure 6

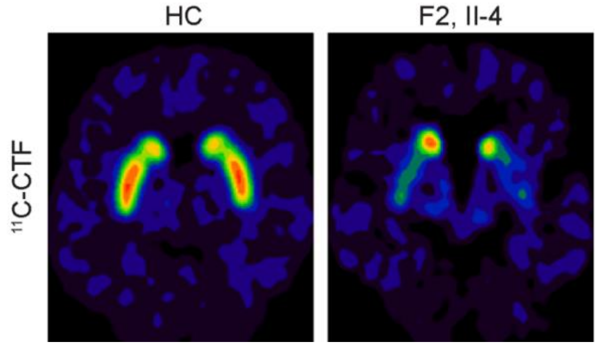


**Fig. S6 PET imaging of PD patient with *DAGLB* mutation.** Representative axial PET images of $^{11}$C-2β-carbomethoxy-3β-(4-fluorophenyl) tropane ($^{11}$C-CFT) uptake in an affected member (Family 2) and healthy control (HC), showing a graded and asymmetrical reduction in dopamine transporter binding ($^{11}$C-CFT) in the putamen.

#### Supplementary Figure 7

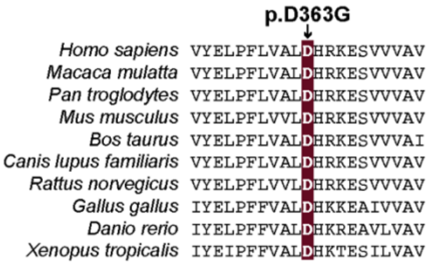


**Fig. S7 The conserved p.D363 residue in the catalytic domain of DAGLB across different species.**

#### Supplementary Figure 8

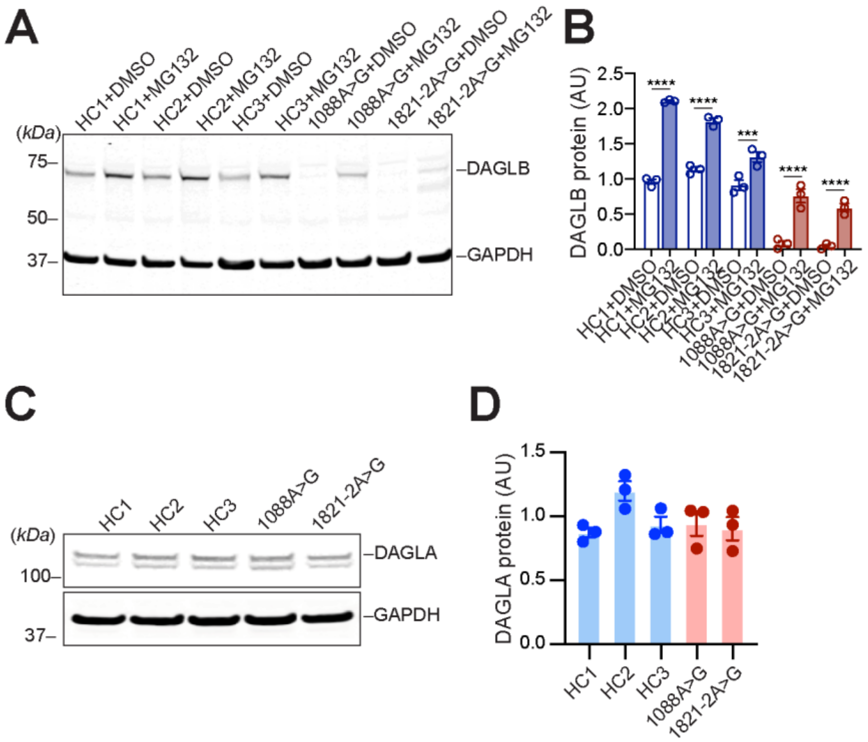


**Fig. S8 Proteasome inhibitor MG132 increased the levels of DAGLB protein in patient-derived fibroblasts. (A)** Representative western blot and **(B)** quantification of protein levels of DAGLB in patient-derived fibroblast cells treated with vehicle or proteasome inhibitor MG132. **(C)** Representative western blot and quantification (**D**) of DAGLA protein levels in patient-derived fibroblast cells. All experiments were performed three times, blinded to genotype (data represent mean ± SEM). Data were normalized to glyceraldehyde 3-phosphate dehydrogenase (GAPDH) protein. 1-way ANOVA with Sidak's multiple comparison test, ***p=0.0002, ****p<0.0001.

#### Supplementary Figure 9

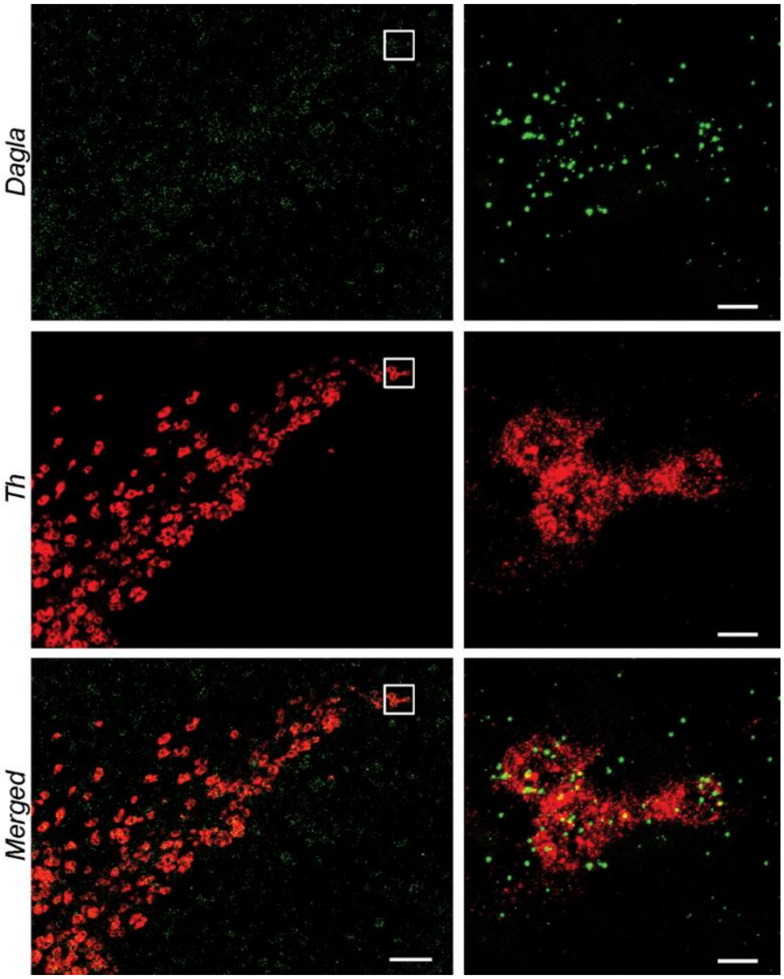


**Fig. S9 RNAscope *in situ* hybridization of *Dagla* and *Th* in mouse midbrain sections.** Right panels highlight the boxed areas in the left panels. Scale bars: 100 μm (left) and 20 μm (right).

#### Supplementary Figure 10

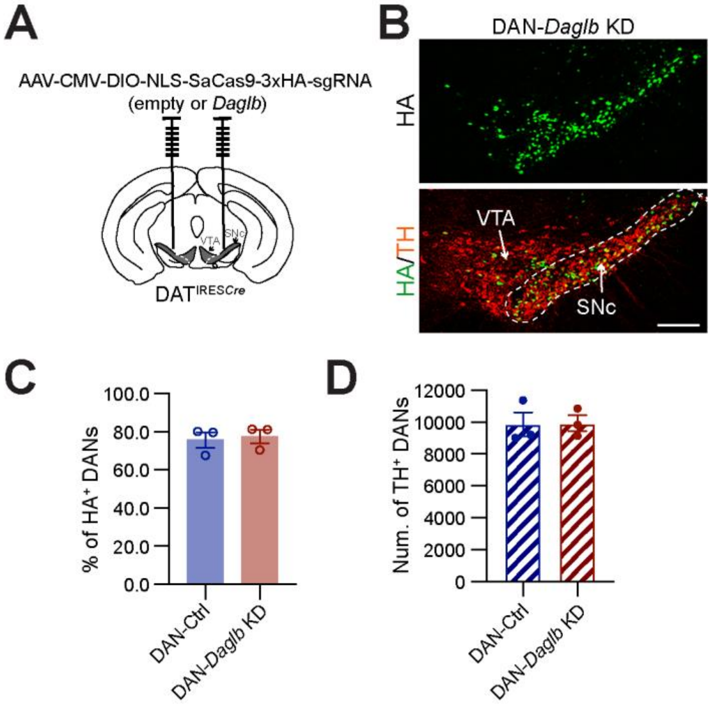


**Fig. S10 Pathological characterization of DAN-*Daglb* KD mice. (A)** Bilateral stereotactic injection of AAV-control and AAV-*Daglb* KD vectors in the SNc of *Dat*$^{\mathrm{IREScre}}$ mice. **(B)** Representative images of HA-SaCas9 (green) and TH (red) staining. Scale bar: 200µm. **(C)** Percentages of HA-SaCas9-postive nigral DANs in the control and KD mice 4-5 months after AAV injection. N=3 per genotype, unpaired t test, p=0.75. **(D)** Numbers of TH-positive nigral DANs in the control and KD mice 12 months after AAV injection. N=3 per genotype, unpaired t test, p=0.94. All experiments were performed blinded to genotype. Data represent mean ± SEM.

#### Supplementary Figure 11

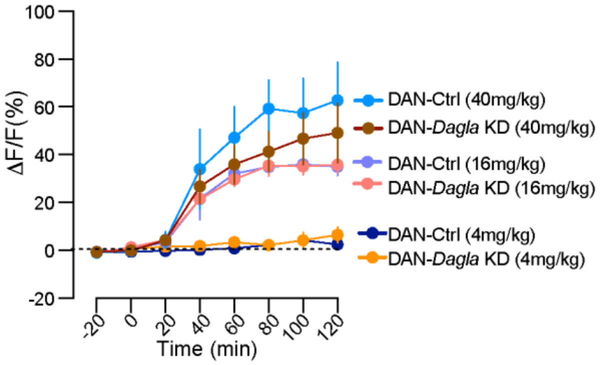


**Fig. S11 DAGLA does not contribute to the major 2-AG synthesis in nigral DANs.** Time course of eCB2.0 signals in the SN of DAN-control and DAN-*Dagla* KD mice before and after treated with JZL184 at 4, 16, and 40 mg/kg. N=4 mice per genotype. Data were presented as mean ± SEM.

#### Supplementary Figure 12

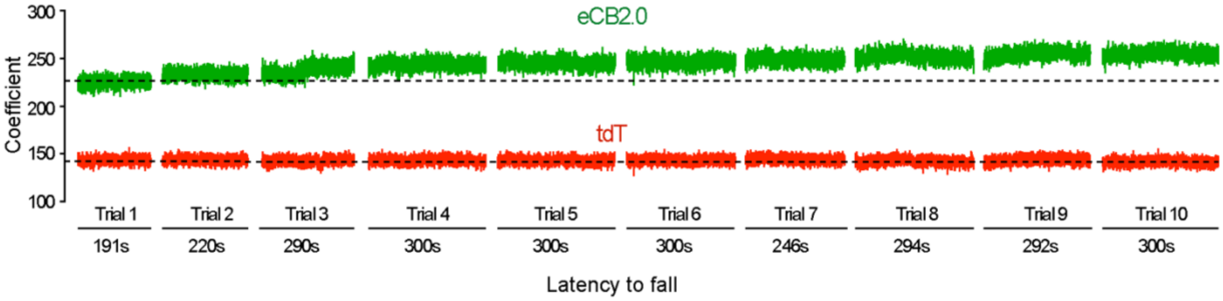


**Fig. S12 Progressive increase of 2-AG signals during rotarod motor skill learning.** Representative raw data of eCB2.0 (green) and tdT (red) signals in the SNr of a mouse performing the 10-trial rotarod motor learning test on day 2 of the 6-day training paradigm.

#### Supplementary Figure 13

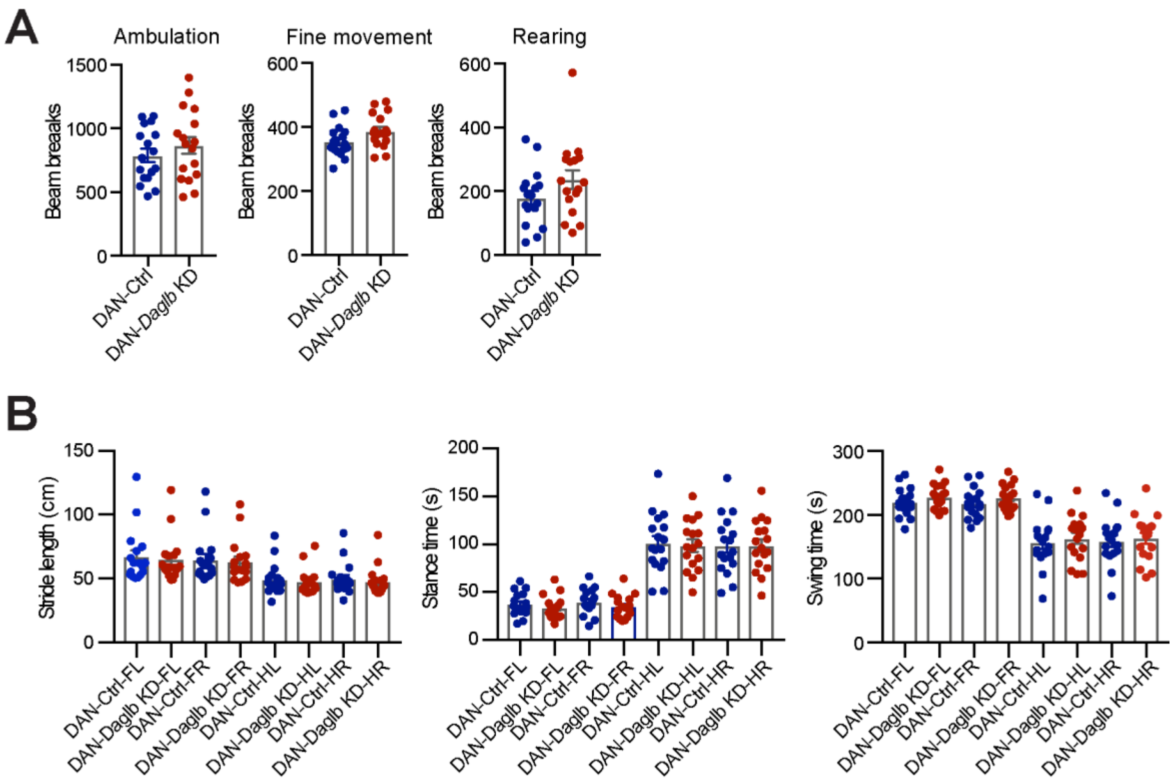


**Fig. S13 Normal locomotion and gait properties of DAN-*Daglb* KD mice. (A)** Open-field tests show comparable travel distance (p=0.23), ambulatory (p=0.30), fine movement (p=0.06), and rearing (p=0.13) between 4-5-month-old *Daglb* control (n=17) and KD (n=17) mice. Data represent mean ± SEM, unpaired t test. **(B)** Gait analyses depict comparable stride length [front left limb (FL): p=0.76, front right limb (FR): p=0.77, hind left limb (HL): p=0.68, hind right limb (HR): p=0.58], stance time (FL: p=0.38, FR: p=0.26, HL: p=0.79, HR: p=0.98) and swing time (FL: p=0.29, FR: p=0.24, HL: p=0.60, HR: p=0.73) of 5-month-old *Daglb* control (n=17) and KD (n=17) mice. Data represent mean ± SEM, unpaired *t* test.

### Supplementary Figure 14

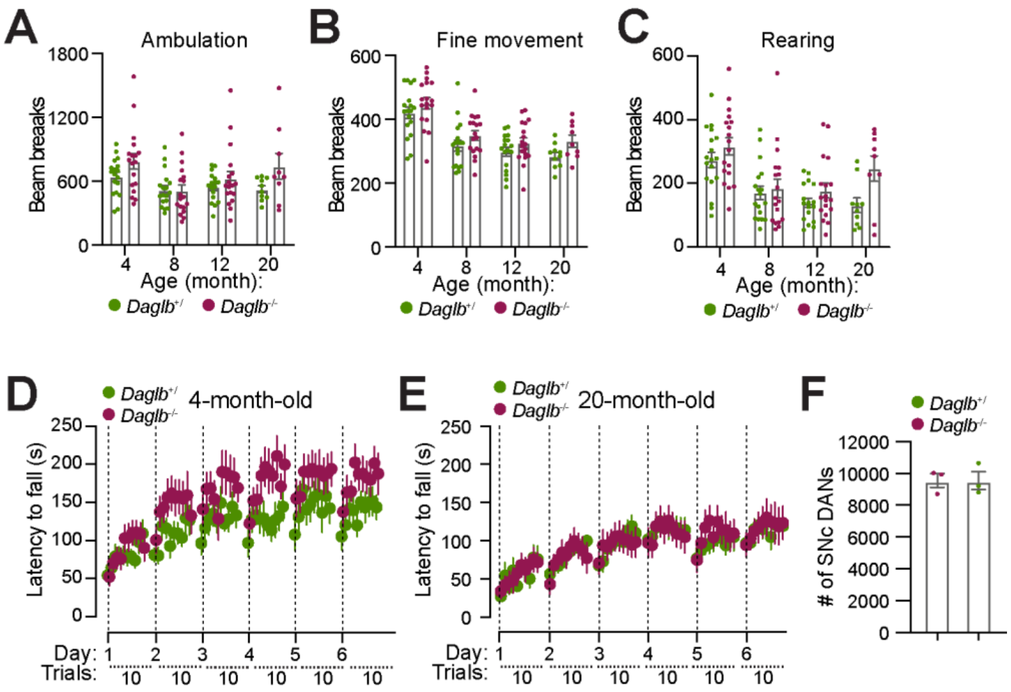


**Fig. S14 Behavioral and neuropathological characterization of *Daglb* germline KO mice.** **(A-C)** Open-field tests show comparable ambulatory (**A**), fine movement (**B**), and rearing (**C**) between the control wild-type and heterozygous *Daglb* KO mice (*Daglb*$^{+/}$, n=10 to 18) and homozygous *Daglb* KO (*Daglb*$^{-/-}$, n=9-18) mice. Data represent mean ± SEM, unpaired t test. (**D-E**) Rotarod motor learning tests of *Daglb*$^{+/}$ and *Daglb*$^{-/-}$ mice at 4 (**D**) and 20 (**E**) months of age. Data represent mean ± SEM. (**F**) Numbers of TH-positive nigral DANs in 20-month-old *Daglb*$^{+/+}$ and *Daglb*$^{-/-}$ mice. N=3 per genotype. Data represent mean ± SEM. Unpaired *t* test, p=0.9989.
